# Chromatin association promotes UBR5-mediated degradation of Rb

**DOI:** 10.64898/2026.04.16.719064

**Authors:** Shuyuan Zhang, Michael C Lanz, Joshua Konschnik, Jan M Skotheim

**Affiliations:** Department of Biological Chemistry and Pharmacology, College of Medicine, The Ohio State University, Columbus, OH 43210, USA; Department of Biology, Stanford University, Stanford, CA 94305, USA; Chan-Zuckerberg Biohub, Stanford, CA 94305, USA

## Abstract

The retinoblastoma protein Rb is a cell cycle inhibitor that plays a central role in regulating the G1/S cell cycle transition. Un-/hypo-phosphorylated Rb suppresses E2F transcription activity by binding to E2F/DP dimers and recruiting chromatin remodelers to prevent cells from entering S phase. For cells to progress through the G1/S transition, Rb is inactivated by two mechanisms: the “classic” pathway of Rb hyperphosphorylation by Cyclin-CDK complexes, and a recently identified “degradation” mechanism driven by the E3 ubiquitin ligase UBR5. These two pathways are interconnected, as only the un-/hypo-phosphorylated Rb can be degraded, and the hyper-phosphorylated Rb is stabilized to promote its reaccumulation in preparation for the next cell division cycle. However, the molecular basis for how Rb is stabilized upon phosphorylation remains unclear. In this study, we found that UBR5 preferentially targets chromatin-associated proteins for degradation. Since Rb’s chromatin association is modulated by its phosphorylation, we hypothesized that phosphorylation may affect Rb stability by altering its chromatin association. To test this, we constructed a series of un-phosphorylatable Rb variants with graded reductions in chromatin association. Consistent with our hypothesis, we observed a strong correlation between an Rb variant’s chromatin association and its half-life. Fusing these Rb variants to histone H1 increased chromatin association to similar levels and equalized their protein half-lives. Taken together, these findings show how phosphorylation stabilizes Rb by promoting its dissociation from chromatin. This provides a striking example for how sub-organellar protein localization may be used to regulate stability.

## Introduction

The retinoblastoma protein Rb is a central cell cycle regulator that inhibits the G1/S transition. To do this, Rb prevents premature entry into S phase by suppressing the activity of E2F transcription factors through both a direct protein-protein interaction and the recruitment of chromatin remodeling proteins^1,2^. To bind the E2F-DP heterodimer, Rb utilizes two distinct domains ^3^. First, the conserved pocket domain of Rb binds the transactivation domain (TD) of the E2F protein^3^, where part of the E2F TD inserts into a groove on the surface of the Rb Pocket.

Second, the C-terminal domain of Rb (RbC) binds to the Marked Box and Coiled-Coil domains of the E2F-DP heterodimer ^2–4^, which anchors Rb and allows it to effectively suppress cell cycle progression. Rb also serves as a scaffold to recruit chromatin remodelers using a single, low-affinity docking site known as the LxCxE cleft that is located on the pocket domain ^5–7^. The recruited chromatin remodelers such as HDAC1, ARID4A, and SUV39H1, can epigenetically modify the nucleosomes, which further suppresses the transcription of E2F-target genes.

The function of Rb can be modulated by its phosphorylation state through two distinct mechanisms. First, phosphorylation of Rb by Cyclin-CDK complexes induces structural changes that disrupt its interactions with E2F and many chromatin-binding partners^1,8^. Second, Rb stability is regulated in a phosphorylation-dependent manner: un-/hypo-phosphorylated Rb is targeted for UBR5 mediated ubiquitination and degradation, whereas phosphorylation stabilizes the protein^9,10^. During early G1 phase, when Rb is un- or hypo-phosphorylated and inhibits E2F^11,12^, it is targeted for degradation by UBR5 to progressively decrease its concentration to promote the G1/S transition^10^. As the cell approaches S phase, Cyclin-CDK complexes become increasingly active and hyper-phosphorylate Rb, disrupting its interactions with E2F and chromatin-binding partners^1,8^. The hyper-phosphorylated Rb is stabilized, leading to an increase of Rb protein concentration during S/G2 phase that resets the regulatory system for the next cell cycle. Thus, Rb phosphorylation functions as a switch that regulates both Rb activity and protein stability.

The molecular mechanisms by which Rb phosphorylation disrupts its interactions with E2F and chromatin remodelers have been studied extensively^1–5,8,11,13–15^, revealing how different Rb phosphorylation sites target specific interaction interfaces on Rb. For example, T373 phosphorylation both inhibits E2F binding and occludes its LxCxE-binding cleft, whereas S608 and S612 phosphorylation selectively disrupts the E2F “silencing” interface. Phosphorylation of S788 and S795, which are located within the C-terminal region of Rb, regulates the high-affinity binding of the Rb C-core domain to E2F^3,11,14^.

While we know a signification amount about how Rb phosphorylation modulates its conformation and E2F interactions, the molecular basis by which Rb phosphorylation prevents its degradation was not understood (Fig. 1A). To address this question, we analyzed features of putative UBR5 substrates and found that UBR5 preferentially targets chromatin-associated proteins. Because the chromatin association of Rb is modulated by its phosphorylation, we hypothesized that Rb’s stability is, at least in part, controlled by its chromatin association. To test this model, we generated a series of un-phosphorylatable Rb variants with various degrees of chromatin association and observed a strong inverse correlation between chromatin association and protein half-life. Moreover, fusing these Rb variants to histone H1 restored their chromatin association to comparable levels and also equalized their protein half-lives. Collectively, these results suggest that Rb’s phosphorylation-dependent degradation by UBR5 is mediated by its chromatin association, providing new molecular insight into the mechanisms governing Rb protein stability at the start of the cell division cycle.

**Figure 1.**
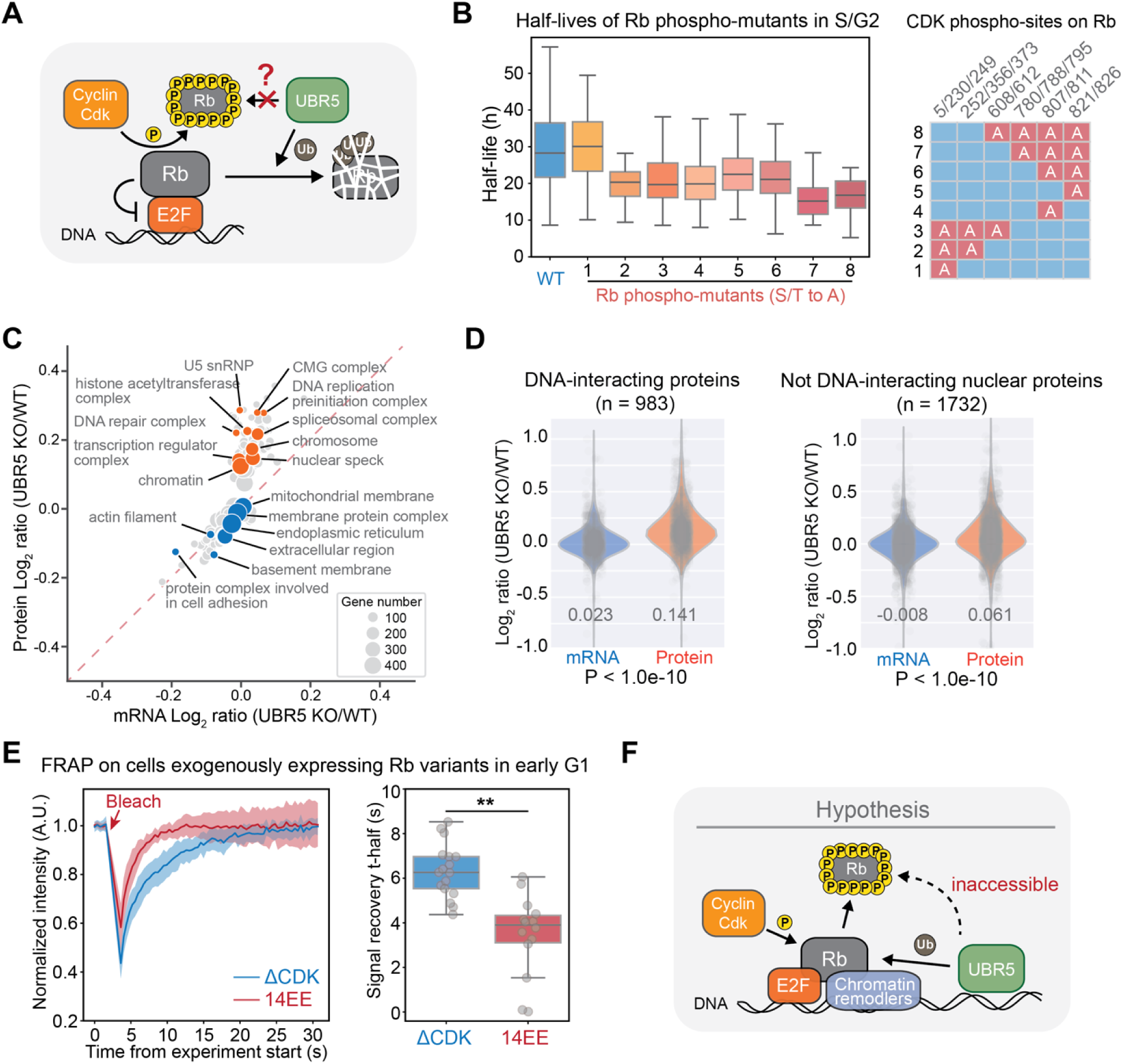
UBR5 targets chromatin-associated proteins, and phosphorylation modulates Rb chromatin association. **A.** Model of phosphorylation-dependent Rb degradation. Un-/hypo-phosphorylated Rb is degraded via UBR5, whereas hyper-phosphorylated Rb is stabilized. **B.** Left panel shows the half-life distributions for the indicated Rb phospho-mutants (S/T to A) in S/G2 phase. Data are from our previous study ^10^. Right panel shows the phosphorylation sites mutated to alanine in each Rb variant. **C.** Annotation enrichment analysis using the mRNA and protein Log_2_ ratio (UBR5 KO/WT) calculated from the mRNA-seq and quantitative proteomics experiments on *UBR5 KO* and *UBR5 WT* cells. Each dot represents a significantly enriched GO Cellular Component annotation group (Benj. Hoch. FDR < 0.02). Dot size reflects the number of proteins in the group, and dot position is determined by the mean Log_2_(UBR5 KO/WT) ratio of genes within the group. The red dotted line denotes when mRNA and protein fold-changes are equivalent. Representative terms are highlighted in orange (upregulated) and blue (unchanged/downregulated). **D.** The Log_2_ ratios (UBR5 KO/WT) in mRNA and protein levels for DNA-interacting protein (left) and Not DNA-interacting nuclear proteins (right). The DNA-interacting genes are defined in ^29^, and the Not DNA-interacting nuclear proteins are defined using the data from the same study^29^, where the proteins are classified as “Nuclear” but not “DNA-interacting”. Each dot represents one gene, and the total number of genes is indicated in the panels. The Log_2_ (UBR5 KO/WT) values and P-value comparing protein and mRNA concentration changes is indicated on the plot. **E.** Left panel shows the mean normalized FRAP recovery curve from HMEC cells expressing Clover-3xFlag-RbΔCDK or Clover-3xFlag-Rb14EE. We examine only cells in early- to mid-G1 as indicated by a CDK activity sensor. The bleach time point is indicated and the shaded area denotes the standard deviation. Right panel shows the distribution of FRAP signal recovery half-time (t-half) for RbΔCDK and Rb14EE. Box plot shows the 5^th^, 25^th^, median, 75^th^, and 95^th^ percentiles. Each dot represents one measured cell. ** denotes *P*<0.01. **F.** Model schematic: UBR5 preferentially targets chromatin-associated Rb for degradation. Hyperphosphorylation promotes Rb dissociation from chromatin making it inaccessible to UBR5 and thereby stabilizing Rb.

## Results

### Rb phosphorylation modulates its chromatin association

In our previous study^10^, we discovered that un-/hypo-phosphorylated Rb is targeted for degradation by the E3 ligase UBR5, whereas Rb hyper-phosphorylation by Cyclin-CDK complexes stabilizes the protein. However, how Rb phosphorylation enables protein stabilization remained unclear (Fig. 1A). A straightforward hypothesis is that, specific Rb phosphorylation sites induce structural changes in Rb that alter its interaction with UBR5. That phosphorylation can alter the association with a particular E3 Ub ligase is common, as in the case of p27Kip1^16^ and Cyclin E^17^, whose phosphorylation promotes their association with SCF. To test this possibility, we re-analyzed the half-lives of a panel of Rb phospho-deficient mutants (S/T to A) in S/G2 phase, as well as phosphomimetic mutants (S/TP to EE) in early G1 phase. We found that, except the 3 N-terminal sites, all remaining 12 sites contributed to Rb protein stability, with no single, double, or triple site combination exhibiting a dominant effect (Fig. 1B; Fig. S1A). Moreover, we observed an additive effect of phosphorylation, such that increasing the number of phosphomimetic sites progressively enhanced Rb stability (Fig. S1A). These results suggest that, rather than acting as a binary on/off switch through a small number of critical sites, Rb phosphorylation induces graded changes that cumulatively promote protein stabilization.

Given that most of Rb’s phosphorylation sites contribute to its stability, it is possible that the negative charge introduced by phosphorylation might underlie the stabilization. To test this, we added a glutamic acid enriched (E-rich) tail to the C-terminus of an un-phosphorylatable Rb variant (RbΔCDK, lacking all CDK phosphorylation sites) (Fig. S1B). We engineered an E-rich tail that contains 30 E residues in order to equal the net negative charge of hyper-phosphorylated Rb (see sequence in Methods). A glutamine-rich (Q-rich) tail was included as a neutral charge control. Using live-cell imaging to measure protein half-lives in early G1 phase (as in our previous study ^10^), we observed that the E-tail RbΔCDK variant showed an increased protein half-life in early G1 phase compared to RbΔCDK. However, the Q-tail variant was similarly stabilized (Fig. S1C), suggesting that negative charge alone cannot account for the phosphorylation-dependent stabilization of Rb.

Since Rb is one of a set of UBR5 substrates, we thought that identifying common features of this set of substrates may help elucidate how phosphorylation stabilizes Rb. To identify these common features, we performed RNA-Seq and quantitative proteomics using *UBR5 WT* and *KO* cells generated in our previous study^10^ (Fig. S2A)(Supplementary table 1, 2). In total, 6413 genes had both non-zero RNA-seq values and >=5 uniquely identified peptides in the proteomics dataset (Fig. S2B). Two-dimensional annotation enrichment analysis revealed that genes whose protein concentrations were either unchanged or decreased in the *UBR5 KO* cells had concordant mRNA changes. In contrast, genes whose proteins increased in concentration in *UBR5 KO* cells had comparatively little changes to their mRNA levels (Fig. 1C). GO cellular component terms indicated that the upregulated protein groups, containing the UBR5 targets, were enriched for nuclear processes, while the unchanged or down-regulated groups were largely cytoplasmic or associated with cytoplasmic organelles (Fig. 1C). This result suggests that nuclear proteins might be preferentially targeted by UBR5, because they exhibit more pronounced changes at the post-transcriptional level. To test this, we took the set of proteins that were significantly altered at the protein level (FDR < 0.05) but not at the mRNA level (FDR > 0.05), defining these as “post-transcriptionally regulated genes” (n = 1177) in *UBR5 KO* cells (Fig. S2C). Gene Ontology (GO) annotations for this gene set were analyzed using SubcellulaRVis^18^ to determine enriched subcellular compartments. This revealed that the nucleus was the only significantly enriched compartment (Fig. S2D). In addition, we classified proteins into distinct categories based on their subcellular location and found that, while the proteins localized to the cytoplasm or cytoplasmic organelles showed minimal changes at both the mRNA and protein levels, the proteins localized to the nucleus and nucleolus exhibited significantly increased protein concentrations relative to their mRNA concentrations (Fig. S3A-E). These observations indicate that UBR5 predominantly targets nuclear proteins.

Our identification of UBR5 targets as generally nuclear is consistent with previous studies showing that UBR5 substrates participate in transcription^10,19–24^, DNA damage response^25–27^, and mitosis^28^, all of which are chromatin-associated processes. Moreover, ChIP-seq experiments revealed that UBR5 associates with numerous chromatin loci^22^, suggesting that it resides in close proximity to chromatin and may thereby target chromatin-associated proteins. To test whether UBR5 preferentially targets this protein class, we compared changes in mRNA and protein concentrations for DNA-interacting proteins identified in a protein-DNA photo-crosslinking study^29^. Protein concentrations for this group were globally increased in *UBR5 KO* cells, whereas the mRNA concentrations remained largely unchanged, resulting in significant differences between mRNA and protein fold changes (Fig. 1D). By contrast, among nuclear proteins that were not classified as DNA-interacting, protein concentrations were not increased to the same extent as for DNA-interacting proteins (Fig. 1D, Log_2_ ratio, 0.141 vs 0.061), although the difference between mRNA and protein fold changes remained significant. Using data from the same study^29^, we also analyzed proteins identified by ChEP (Chromatin Enrichment for Proteomics), which enriches for chromatin-associated proteins, and compared them with nuclear proteins not identified by ChEP. While mRNA concentrations remained unchanged for both groups, ChEP-enriched proteins showed a greater increase in protein concentration than non-ChEP nuclear proteins (Fig. S3F, Log_2_ ratio, 0.102 vs 0.028). Collectively, these results suggest that UBR5 preferentially targets chromatin-associated proteins for degradation.

That UBR5 preferentially targets chromatin-associated proteins suggested that Rb’s chromatin association being restricted to G1 might underlie its stability dynamics. In early- to mid-G1 phase, un-/hypo-phosphorylated Rb associates with chromatin via binding to E2F and chromatin remodelers^30,31^. Rb’s hyper-phosphorylation in late G1 promotes its dissociation from E2F^3,4,8,14,15,32,33^ and thus alters its chromatin association^34,35^. To confirm that phosphorylation alters Rb’s chromatin association, we used Fluorescence Recovery After Photobleaching (FRAP) to measure the signal recovery kinetics of unphosphorylated Rb (Clover-Flag-RbΔCDK) and phosphomimetic Rb (Clover-Flag-Rb14EE) (Fig. S4A). The negative controls, Clover and Clover-NLS recovered very rapidly after photo-bleaching, as compared to Clover-Flag-RbΔCDK (Fig. S4B). We note that Rb14EE was used instead of the more stable variant Rb13EE, because the fluorescence intensity of Rb13EE was substantially higher than RbΔCDK, preventing the use of identical laser settings. Importantly, Rb14EE and Rb13EE exhibited indistinguishable FRAP kinetics (Fig. S4C) and are therefore interchangeable for assessing chromatin association. Rb14EE exhibited a significantly faster recovery rate than RbΔCDK (Fig. 1E), indicating that phosphorylation increases Rb mobility and reduces its chromatin association. These data are consistent with a model in which phosphorylation of Rb disrupts its chromatin association to increase its mobility in the nucleus.

Since the addition of phosphomimetic sites affected the mobility of an exogenously expressed Rb protein in the nucleus, likely through its chromatin association, we sought to test if the phosphorylation of endogenous Rb in G1 phase impacted its mobility. To do this, we examined an HMEC cell line containing an endogenously Clover-tagged Rb and a CDK activity sensor^36^. The presence of the CDK activity sensor allowed us to classify cells into CDK-low and CDK-high states, which correlates well with hypo- and hyper-phosphorylated Rb, respectively^37^.

Consistent with our findings obtained using exogenously expressed constructs, endogenous Rb recovered from photobleaching significantly slower in CDK-low cells compared to CDK-high cells (Fig. S4D). Together, these results support a model in which Rb phosphorylation decreases its chromatin association, thereby reducing its likelihood of being targeted by UBR5 and ultimately stabilizing the Rb protein (Fig. 1F).

### Generating Rb variants with reduced protein interactions

If Rb’s chromatin association controls its stability, then disrupting Rb’s chromatin association should stabilize the protein. To test this, we first needed to generate Rb variants with reduced chromatin association. Rb associates with chromatin through interactions with the E2F-DP complex and chromatin remodelers ^30,31^. We therefore introduced mutations into Rb to disrupt these interactions. The selection of residues for mutation was guided by previously published structures and mutational results^4,5,38,39^ (Fig. S5B-D). Briefly, Rb interacts with the E2F-DP heterodimer through two interfaces^3^. The first interface is located within the Rb pocket domain and binds the transactivation domain (TD) of E2F^3^. To disrupt this interaction, we introduced the previously described Rb ΔG mutation^38,40^ (annotated as “dG”) (Fig. 2A, Fig. S5A, B). The second interface involves the Rb C-terminal domain (RbC), which binds the Marked Box and Coiled-Coil domains of the E2F-DP heterodimer^2–4^. Based on the previously solved structure (PDB: 2AZE)^4^, we performed alanine substitutions at seven residues (annotated as “C7A”) to disrupt this interaction (Fig. 2A, Fig. S5A, C). In addition, Rb recruits chromatin remodelers through the LxCxE-binding cleft located within the pocket domain^5–7^. To disrupt this interaction, we mutated three residues (annotated as “LCE”)^5,39,41^(Fig. 2A, Fig. S5A, D). To eliminate the confounding effect of phosphorylation, all mutations were introduced into un-phosphorylatable Rb protein (RbΔCDK), ensuring that none of the variants could be phosphorylated by CDKs. Consistent with the known roles of these interactions, RbΔCDK expression induced G1 arrest (Fig. S6A), whereas all mutant RbΔCDK variants exhibited a reduced cell cycle arrest (Fig. S6A). Notably, the effects of different mutation types are additive so that the variants with combinations of mutations displayed a higher proportion of cells in S/G2 phases (Fig. S6A).

**Figure 2.**
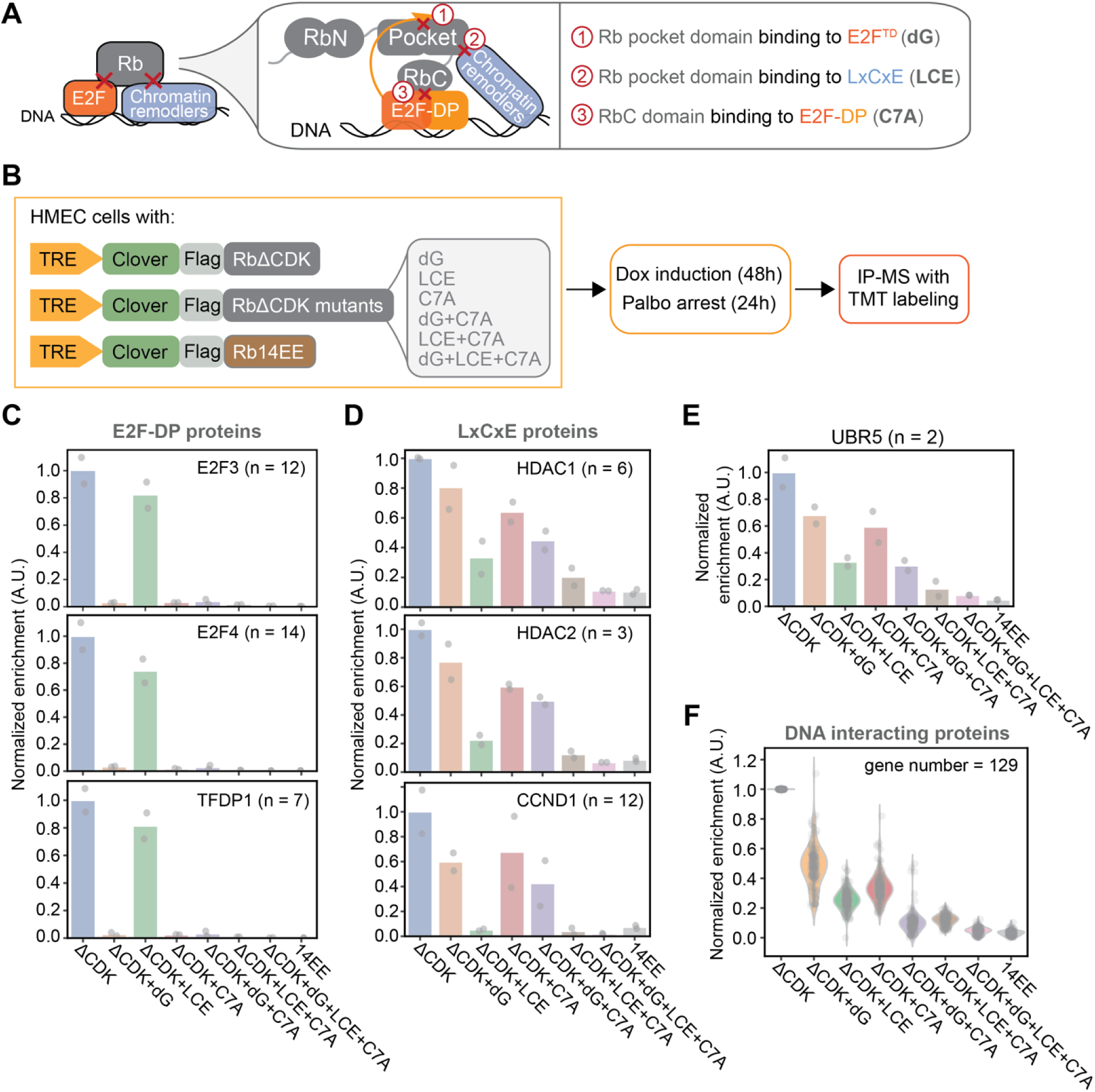
Generation and validation of Rb variants that selectively disrupt Rb interactions. **A.** Schematic of the three major Rb interaction surfaces that bind the E2F-DP complex and chromatin remodelers, along with abbreviated names for the corresponding sets of mutations. **B.** Workflow for IP-MS experiment used to define the interactomes of different Rb variants. **C.** The normalized enrichment of E2F-DP associated proteins identified by IP-MS across different Rb variants. Points represent two replicate IP-MS experiments. n denotes the number of unique peptides detected for each protein. **D.** The normalized enrichment of LxCxE containing proteins interacting with the indicated Rb variants identified by IP-MS. Points represent two replicate IP-MS experiments. n denotes the number of unique peptides detected for each protein. **E.** The normalized enrichment of UBR5 across different Rb variants. Two unique peptides were detected for UBR5 (n = 2). Each dot represents one replicate IP-MS experiment. **F.** The normalized enrichment of DNA-interacting proteins identified by IP-MS across different Rb variants. Each point is the average of two replicate IP-MS experiments for a different DNA-interacting protein. The DNA-interacting proteins are defined in ^29^, and a total of 129 DNA-interacting proteins were identified in this IP-MS experiment.

To verify that the three sets of Rb mutations, namely, dG, C7A, and LCE, disrupted the intended interactions, we performed immunoprecipitation-mass spectrometry (IP-MS) to quantitatively profile the interactomes of the different RbΔCDK variants (Fig. 2B; Supplementary table 3). Briefly, HMEC cells were stably transduced with a doxycycline (Dox)-inducible system in which cells conditionally express Clover-3xFlag-tagged Rb variants upon Dox treatment (1µg/mL). After 48h of Dox treatment, we harvested cells for immunoprecipitation and mass-spectrometry. Phosphomimetic Rb (Rb14EE) was included as a control, and cells with no Rb vector served as a negative control to remove nonspecific interactions. Since cells expressing different variants have different cell cycle dynamics (Fig. S6A), we sought to ensure that interactomes were compared within the same cell cycle phase. To do this, cells were synchronized in G1 through treatment with Palbociclib (1μM) for 24 hours before harvesting (Fig. 2B). We then used tandem mass tag (TMT) labeling to compare the Rb interactome across protein variants. In total, 2472 proteins were identified. The interacting proteins were defined as those showing a >= 2-fold higher proportion in at least one Rb bait sample compared with the empty mock sample, and identified by >=3 unique peptides (n = 924 Rb interacting proteins). Because different Rb variants were expressed at different levels (Fig. S7A), the proportion of each interacting protein was normalized to the corresponding bait protein to generate a normalized enrichment score, with the RbΔCDK set to 1. To validate the impact of each mutation on the Rb interactome, we examined the normalized enrichment of E2F-DP proteins and known LxCxE-containing proteins and compared these across the different Rb variants. As predicted, Rb variants harboring dG and/or C7A mutations showed a markedly reduced interaction with E2F-DP proteins, while still interacting with LxCxE-containing proteins. Conversely, Rb variants carrying LCE mutations exhibited substantially reduced binding to LxCxE-containing proteins while maintaining interactions with E2F-DP proteins (Fig. 2C, D). These results confirm that the introduced Rb mutations selectively disrupted their intended interaction interfaces.

Because these Rb variants were designed to alter chromatin association, we next examined their interactions with a defined set of DNA-interacting proteins ^29^. A total of 129 DNA-interacting proteins were identified in our interactome dataset, and all RbΔCDK variants exhibited reduced interactions with this group of proteins (Fig. 2F; Supplementary table 4). Moreover, Rb variants with more extensively disrupted interfaces displayed further reduced associations, and the phosphomimetic Rb14EE showed the lowest level of interaction with DNA-interacting proteins (Fig. 2F). Although only two peptides corresponding to UBR5 were detected in our IP-MS experiment, the interacting pattern between Rb variants and UBR5 closely mirrored that of DNA-interacting proteins (Fig. 2E). Collectively, these data demonstrate that the engineered Rb mutations selectively disrupt their corresponding protein interactions, and these mutations also reduce Rb’s interaction with chromatin-associated proteins.

### Disrupting Rb’s protein interactions decreases its chromatin association

Having generated a series of Rb variants with disrupted interactions with E2F-DP and LxCxE-containing proteins, we next examined whether these alterations affect Rb chromatin association. To do this, we first used a fluorescence recovery after photobleaching (FRAP) assay to measure the recovery kinetics of these variants, which reflect their degree of chromatin association (Fig. S8A). Rb variants more tightly associated with chromatin would diffuse more slowly and the bleached fluorescence would recover more slowly. Briefly, HMEC cells were stably transduced with a doxycyclin (Dox)-inducible system in which cells conditionally express Clover-3xFlag-tagged RbΔCDK variants upon Dox treatment (1µg/mL). After 48h of Dox treatment, we performed FRAP experiments (Fig. S8A). Since cells expressing different variants have different cell cycle dynamics, with RbΔCDK expressed cells mainly arrested in G1 and in a low CDK-state (Fig. S6A), we needed a way to control for cell cycle phase. To do this, we added a CDK sensor^36^ that allows us to only select cells in a CDK-low G1 state for FRAP measurements. The results showed that all RbΔCDK variants exhibited faster signal recovery compared to RbΔCDK (Fig. 3A, B, Fig. S8B, C), indicating a reduced chromatin association. Notably, variants with more extensively disrupted interfaces showed progressively faster recovery kinetics. The phosphomimetic Rb14EE variant displayed the fastest recovery and lowest chromatin association among all variants (Fig. 3A, B, Fig. S8B, C).

**Figure 3.**
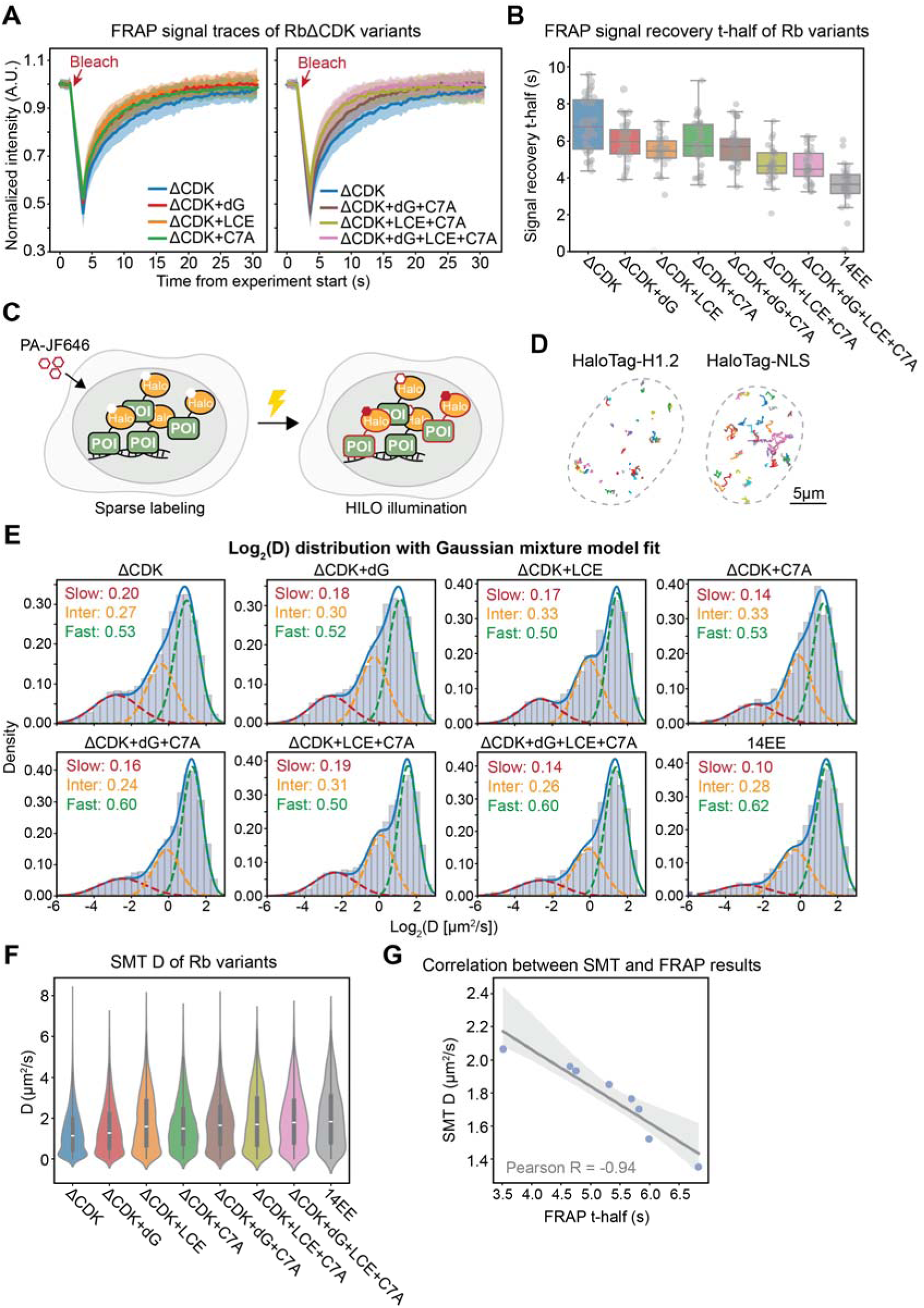
Disrupting Rb interactions with its binding partners decreases its chromatin association. **A.** Mean normalized FRAP recovery curve from HMEC cells expressing Clover-3xFlag-RbΔCDK or the related RbΔCDK interaction-defective variants. To emphasize differences, the left panel compares RbΔCDK to variants carrying a single mutation set, and the right panel compares RbΔCDK to variants carrying two or three mutation sets. The bleach time point is indicated and the shaded areas denote the standard deviation. **B.** Distribution of FRAP signal recovery half-time (t-half) for the indicated Rb variants. Box plot shows the 5^th^, 25^th^, median, 75^th^, and 95^th^ percentiles. Each dot represents one cell. **C.** Schematic of the single molecule tracking (SMT) experiment. Cells expressing HaloTag fused proteins of interest (POI) are labeled with low concentrations of PA-JF646 dye to achieve sparse labeling and then imaged in live cells on a Zeiss Elyra 7 microscope using HILO (Highly Inclined and Laminated Optical sheet) illumination mode. Low activation laser power was used to activate only a subset of molecules, enabling detection of spatially separated single molecule spots for tracking. **D.** Representative single molecule tracks of HaloTag-H1.2 or HaloTag-NLS from two characteristic cells. The dotted lines mark nuclear boundaries. **E.** Probability densities of Log_2_ transformed apparent diffusion coefficients (Log_2_ D) for each Rb variant. The probability densities were fit using a Gaussian mixture model (GMM). The solid blue line shows the overall GMM probability density function (PDF), and dotted curves show the PDFs of the individual mixture components. The fraction of molecules assigned to each diffusive state (Slow, Intermediate, Fast) was estimated from the fitted mixture weights and is indicated on the plot. **F.** Violin plot of the apparent diffusion coefficient (D) across all Rb variants measured from SMT experiments. Inner box plots indicate the first quartile, median, and third quartile. **G.** Pearson correlation between the diffusion coefficients D from SMT and the FRAP recovery half-time (t-half) for the Rb variants. Shaded area denotes the 95% confidence interval of a linear fit.

To validate our chromatin association measurements of different Rb variants, we employed a single molecule tracking assay (SMT; see methods)^42,43^. Using the Piggybac transposon system, we generated stable cell lines harboring a doxycycline (Dox)-inducible cassette expressing HaloTag fused Rb variants (Fig. S9A). Cells were induced with Dox for 48 hours, labeled with low concentrations of PA-JF646 dye, and subjected to single-molecule live-cell imaging using a Zeiss Elyra 7 microscope operated in HILO (Highly Inclined and Laminated Optical sheet) illumination mode (Fig. 3C, Fig. S9A). These cells also harbor the CDK sensor^44^ so that we only selected cells in the CDK-low state for our SMT measurements. The apparent diffusion coefficient (D) of individual Rb molecules was calculated from the stepwise mean-squared displacement (MSD) of their trajectories. The different diffusing modes were identified by fitting the distribution of D using a Gaussian mixture model comprising three fractions^45^. To validate this SMT approach, we generated cell lines expressing either HaloTag-Histone H1.2, a chromatin bound protein, or HaloTag-NLS, which is localized to the nucleus but not tightly bound to chromatin. SMT analysis showed that HaloTag-H1.2 exhibited relatively immobile trajectories, whereas HaloTag-NLS molecules were substantially more mobile (Fig. 3D). Consistent with this observation, HaloTag-H1.2 displayed a much lower D and a larger slow-diffusing fraction compared to HaloTag-NLS (Fig. S9B, C), confirming that our SMT measurements capture the chromatin association states of the tracked molecules.

After establishing the SMT method to examine chromatin association, we then applied this approach to different Rb variants. As expected, RbΔCDK showed the lowest D and the largest slow-diffusing fraction, while all the other variants exhibited increased D values and higher intermediate-to-fast diffusing fractions (Fig. 3E), consistent with our previous FRAP measurements indicating a reduced chromatin association. Moreover, variants with more extensively disrupted interfaces showed further increased D, with the phosphomimetic Rb14EE being the fastest diffusing variant (Fig. 3E, F). Importantly, the results obtained from SMT strongly correlated with those from our FRAP analyses (Fig. 3G; Fig. S9D). Taken together, these complementary approaches demonstrate that disrupting Rb’s interactions with its binding partners decreases its chromatin association, and that increasing the extent of interaction disruption results in progressively weaker chromatin association.

### Rb variants with decreased chromatin association exhibit increased protein stability

After generating a series of un-phosphorylatable Rb variants that display distinct degrees of chromatin association, we sought to measure their half-lives to test our model that Rb chromatin association regulates protein stability. To measure half-lives of these chromatin association variants, we used live-cell imaging as described previously^10^ (Fig. 4A). Briefly, HMEC cells were stably transduced with a doxycyclin (Dox)-inducible system that we used to conditionally express Clover-3xFlag-tagged RbΔCDK variants upon Dox treatment (1µg/mL). After 48h of Dox treatment, we withdrew Dox and monitored the decrease of the Clover fluorescence signal using live cell imaging. Because cells expressing different Rb variants exhibit different cell cycle dynamics, with RbΔCDK-expressing cells predominantly arrested in G1 and exhibiting low CDK activity (Fig. S6A), we also included a CDK sensor^44^ into these cells to enable assessment of protein degradation specifically in CDK-low G1 cells. By fitting the fluorescence decay traces to a simple exponential decay function, we obtained the half-life of each Clover-3xFlag-Rb variant in individual CDK-low G1 cells.

**Figure 4.**
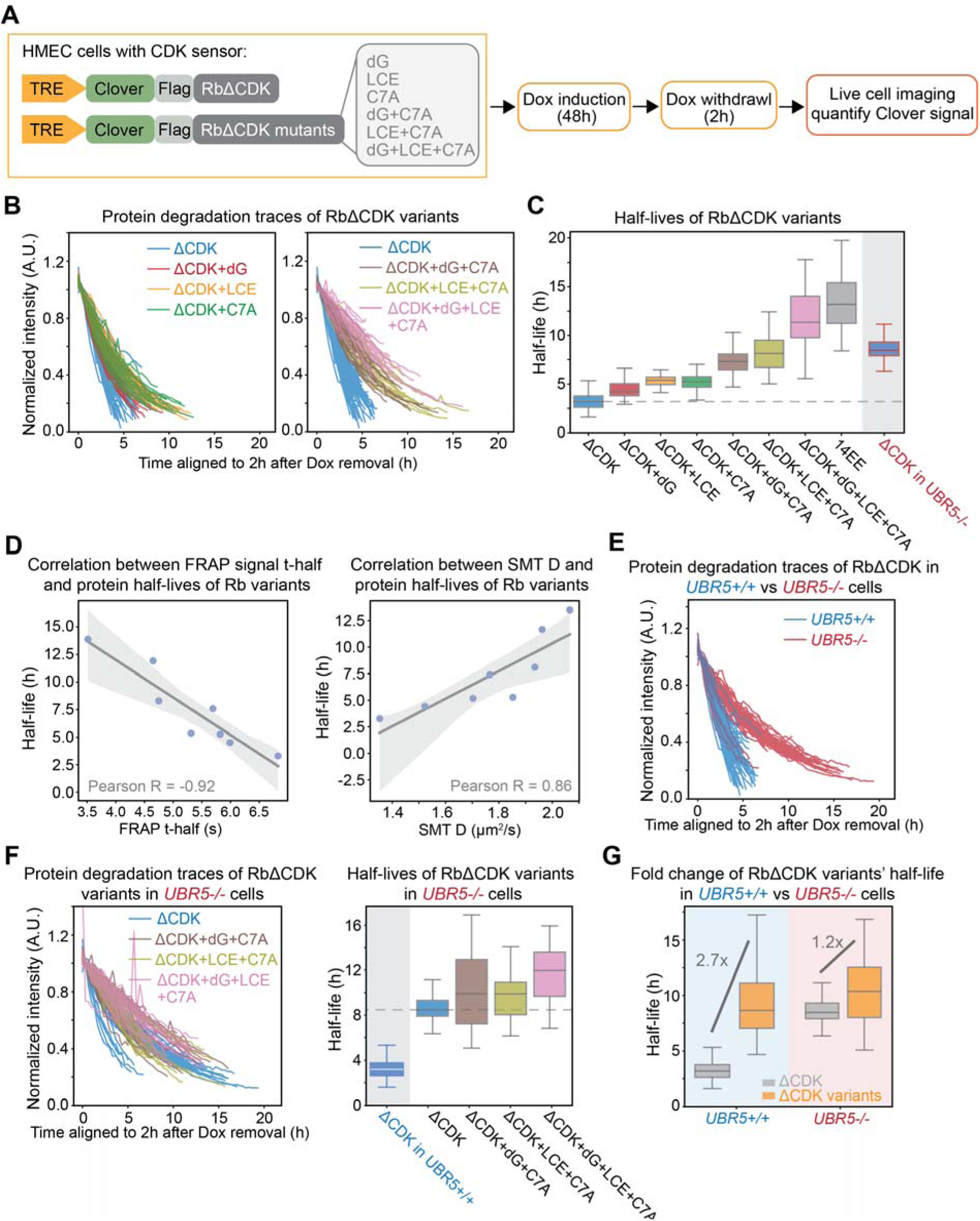
Rb variants with decreased chromatin association are more stable. **A.** Workflow schematic for the live-cell imaging experiment to measure the protein half-lives of Rb variants. **B.** Degradation traces of Clover-3xFlag-Rb variants following Dox withdrawal in wild-type HMEC cells. To highlight differences, the left panel compares RbΔCDK to variants carrying a single mutation set, and the right panel compares RbΔCDK to variants carrying two or three mutation sets. Only the traces in early G1 phase were selected based on a FUCCI cell cycle marker and cell cycle phase duration. **C.** Distribution of half-lives estimated by exponential fitting of the traces in (**B**) (*UBR5+/+*) and (**E**) (*UBR5-/-)*. Half-life data for 14EE is from Figure S1A. Box plot indicates 5^th^, 25^th^, median, 75^th^, and 95^th^ percentiles. **D.** Pearson correlation between the protein half-lives of Rb variants in (**C**) and the FRAP recovery half-time (t-half) (Left panel) or diffusion coefficients D measured from SMT (Right panel). Shaded area denotes the 95% confidence interval. **E**. Degradation traces of Clover-3xFlag-RbΔCDK following Dox removal in *UBR5+/+* (blue) or *UBR5-/-* (red) HMEC cells. Only the traces in early G1 phase were selected, and the classification was based on a FUCCI cell cycle marker and cell cycle phase duration. **F.** Left panel shows the degradation traces of Clover-3xFlag-RbΔCDK and variants carrying two or three mutation sets after Dox withdrawal in *UBR5-/-* cells. Only the traces in early G1 phase were selected, and the classification was based on a FUCCI cell cycle marker and cell cycle phase duration. Right panel shows the distribution of half-lives estimated from exponential fits of the traces shown in the left panel. Box plot indicates 5^th^, 25^th^, median, 75^th^, and 95^th^ percentiles. **G.** Half-life distribution for RbΔCDK (grey) and pooled RbΔCDK variants carrying two or three mutation sets (orange) in *UBR5+/+* and *UBR5-/-* HMEC cells. Variant half-lives were pooled together from dG-C7A, LCE-C7A, and dG-LCE-C7A in (**C**) or (**F**). The fold-changes relative to RbΔCDK are indicated in the plot.

Our half-life measurements revealed that all RbΔCDK variants with disrupted protein interactions exhibited increased half-lives compared to RbΔCDK. The variants harboring more extensively disrupted interfaces displayed the greatest increase in protein stability (Fig. 4B, C). Moreover, there is a strong correlation between the half-lives of Rb variants and their chromatin association as measured by FRAP and SMT (Fig. 4D; Fig. S9E). These findings support our model that reduced chromatin association decreases the likelihood of Rb interacting with UBR5, thereby lowering its rate of being targeted for degradation. To further support this model, we examined the protein stability of these variants in a *UBR5-/-* background. If our model is correct, the half-lives of the RbΔCDK variants with disrupted protein interaction surfaces should be similar to that of RbΔCDK in the absence of UBR5. As expected, while RbΔCDK was markedly stabilized in *UBR5-/-* cells (Fig. 4C, E), the variants with more extensively disrupted interfaces (dG-C7A, LCE-C7A, dG-LCE-C7A) did not show a substantial additional increase in half-life relative to RbΔCDK in *UBR5-/-* cells (Fig. 4F). We further quantified the fold-change in half-life of these RbΔCDK variants (dG-C7A, LCE-C7A, dG-LCE-C7A) relative to RbΔCDK in both *UBR5+/+* and *UBR5-/-* cells. This analysis revealed an approximately 2.7-fold increase in half-life in *UBR5+/+* cells and a modest 1.2-fold increase in *UBR5-/-* cells (Fig. 4G).

Taken together, our results show that reduced chromatin association confers increased protein stability, but only in cells containing UBR5. This supports a model in which Rb’s chromatin association regulates its degradation via UBR5.

### Tethering Rb variants to chromatin prevents their stabilization

If Rb chromatin association regulates its stability, then forcibly tethering Rb variants to chromatin should decrease their stability (Fig. 5A). To directly test this, we artificially increased the chromatin association of the Rb variants with more extensively disrupted interaction interfaces by fusing them to Histone H1 (Fig. 5B). Histone H1.1 exhibits strong chromatin association on its own, and fusion to Rb variants significantly increased their chromatin association, as measured by our FRAP assay (Fig. 5C, D; Fig. S10A-C). We also fused Rb variants to another histone H1 protein, H1.4, and observed a similarly robust increase in chromatin association in the corresponding fusion proteins (Fig. S10D-H). Note that while RbΔCDK alone did not show an immobile fraction in our FRAP assay (Fig. S10C, F), all the Histone H1 fused variants, including Histone H1 itself, exhibited a range of immobile fractions (Fig. S10C, F), which is consistent with previous FRAP measurements of Histone H1^46^.

**Figure 5.**
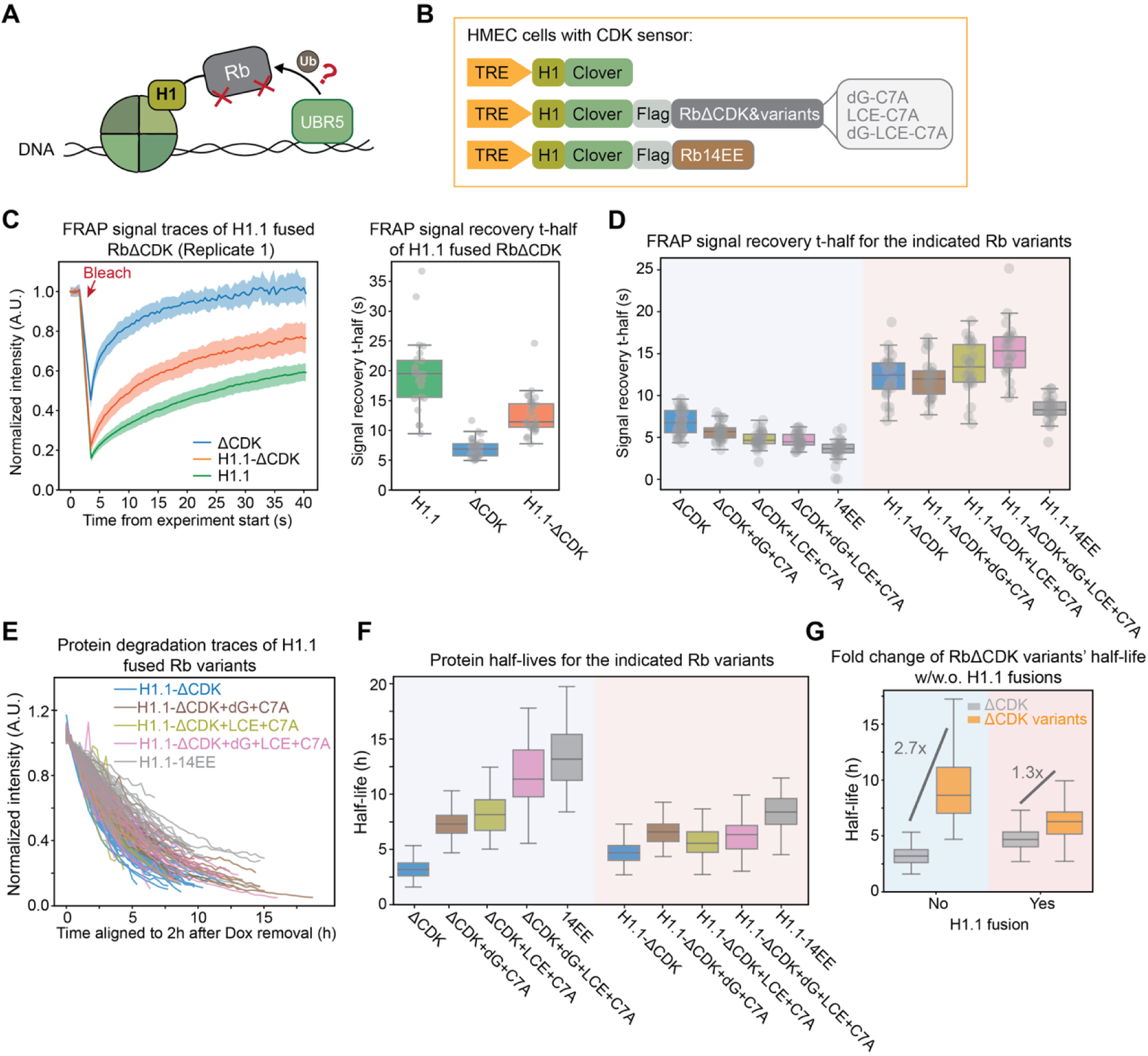
Tethering Rb variants to chromatin prevents their stabilization. **A.** Schematic illustrating the strategy of forcing Rb variants onto chromatin by fusing each variant to histone H1. **B.** Schematic of the engineered cell lines expressing H1-fused Rb variants. **C.** Left panel shows the mean normalized FRAP recovery curves from HMEC cells expressing H1.1-Clover, H1.1-Clover-3xFlag-RbΔCDK, or Clover-3xFlag-RbΔCDK Rb protein variants. The bleach time point is indicated and the shaded area denotes the standard deviation. Right panel shows the distribution of FRAP signal recovery half-time (t-half) for the lines in left panel. Box plot shows the 5^th^, 25^th^, median, 75^th^, and 95^th^ percentiles. Each dot represents one cell. **D.** Distribution of FRAP signal recovery half-time (t-half) for the H1.1-fused Rb variant expressing cell lines (**Fig. S10A-B)** compared with the corresponding Rb variants not fused to H1.1 (Fig. 3B). Box plot shows the 5^th^, 25^th^, median, 75^th^, and 95^th^ percentiles. Each dot represents one cell. **E.** Degradation traces of H1.1-fused Clover-3xFlag-Rb variants following Dox removal. Only the traces in early G1 phase were selected, and the classification was based on a FUCCI cell cycle marker and cell cycle phase duration. **F.** Distribution of half-lives estimated by exponential fits of the traces in (**E**), compared with the corresponding non-H1.1-fused Rb variants (Fig. 4C). Box plot indicates 5^th^, 25^th^, median, 75^th^, and 95^th^ percentiles. **G.** Half-life distributions for RbΔCDK (grey) and pooled RbΔCDK variants carrying two or three mutation sets (orange) with or without H1.1 fusion. Variant half-lives were pooled together from dG-C7A, LCE-C7A, and dG-LCE-C7A in (**F**). The fold-changes relative to RbΔCDK are indicated in the plot.

After establishing that H1 fusion promotes Rb chromatin association, we next measured the protein half-lives of the H1-fused Rb variants using the same live-cell imaging approach and compared them with those of the corresponding unfused Rb variants. To exclude the possibility that fusion to histone H1 induces rapid protein degradation, which can artificially equalize degradation rates across variants, we first measured the half-lives of histone H1.1 and H1.4 alone. Both H1 variants were highly stable, with half-lives ranging from 30 to 40 hours (Fig. S11A, B), substantially longer than those observed for any Rb variants. Thus, any differences in the half-lives of H1-fused Rb variants are unlikely to be due to H1 itself. The half-life measurements of the H1-fused Rb variants showed that H1 fusion prevented these variants from exhibiting significant increases in their half-lives relative to RbΔCDK (Fig. 5E, F; Fig. S11D, E). Notably, even the phosphomimetic Rb14EE displayed a significantly reduced half-life upon H1 fusion (Fig. S11C, G). We further quantified the fold-change in half-life of these RbΔCDK variants (dG-C7A, LCE-C7A, dG-LCE-C7A) relative to RbΔCDK with or without H1-fusion. This analysis revealed an approximately 2.7-fold increase in half-life of non-H1-fused variants and a modest 1.3-fold and 1.1-fold increase for H1.1. and H1.4 H1-fused variants, respectively (Fig. 5G; Fig. S11F). Collectively, these data demonstrate that tethering Rb variants to chromatin via H1 fusion abolishes their stabilization relative to RbΔCDK. This further supports our conclusion that Rb is preferentially degraded by UBR5 when it is associated with chromatin.

## Discussion

Two distinct pathways impinge on Rb to promote the G1/S transition. First, there is the canonical phosphorylation pathway, initiated by Cyclin D-Cdk4/6 complexes and completed by Cyclin E/A-Cdk2 complexes, that inactivates Rb by promoting its dissociation from the activating E2F transcription factors^11,37,47^. Second, there is a non-canonical pathway in which Rb is targeted for degradation by UBR5 during early G1 phase to promote the G1/S transition^9,10^. This degradation of Rb is phosphorylation dependent: Rb is degraded when un-/hypo-phosphorylated and stabilized when hyper-phosphorylated by Cyclin-CDK complexes^9,10^. This stabilization during S/G2/M phases allow its reaccumulation in preparation for the next cell cycle. However, how phosphorylation stabilizes Rb was unclear.

Here, through a mutational analysis, we found that Rb stabilization is unlikely to be driven by a few dominant phospho-sites, but instead reflects the cumulative effect of multisite phosphorylation (Fig. 1B; Fig. S1A). This cumulative effect is unlikely to be driven by a purely charge-mediated mechanism since the addition of the same amount of charge via a glutamic acid tail did not stabilize Rb. Because UBR5 preferentially targets chromatin-associated proteins and because Rb phosphorylation alters its chromatin association (Fig. 1C-E; Fig. S3F, S4D), we proposed that phosphorylation regulated Rb’s stability by changing its chromatin association. To test this model, we generated a series of un-phosphorylatable Rb variants with graded chromatin association by disrupting interactions with the E2F-DP dimer and LxCxE containing proteins. These variants exhibited progressively reduced chromatin association and increased protein half-lives that were well correlated with their chromatin association. Such stabilization of Rb was only observed in *WT* but not *UBR5 KO* cells, indicating that UBR5 was indeed the E3 ligase responsible for the chromatin-dependent degradation of Rb. Moreover, forcing Rb to be more chromatin associated via fusing it to histone H1 promoted its degradation. Together, these findings support a model in which Rb is degraded by UBR5 when associated with chromatin in complexes with E2F-DP and chromatin remodelers. Hyper-phosphorylation in late G1 disrupts these interactions, leading to Rb dissociation from chromatin and protection from UBR5 (Fig. 6).

**Figure 6.**
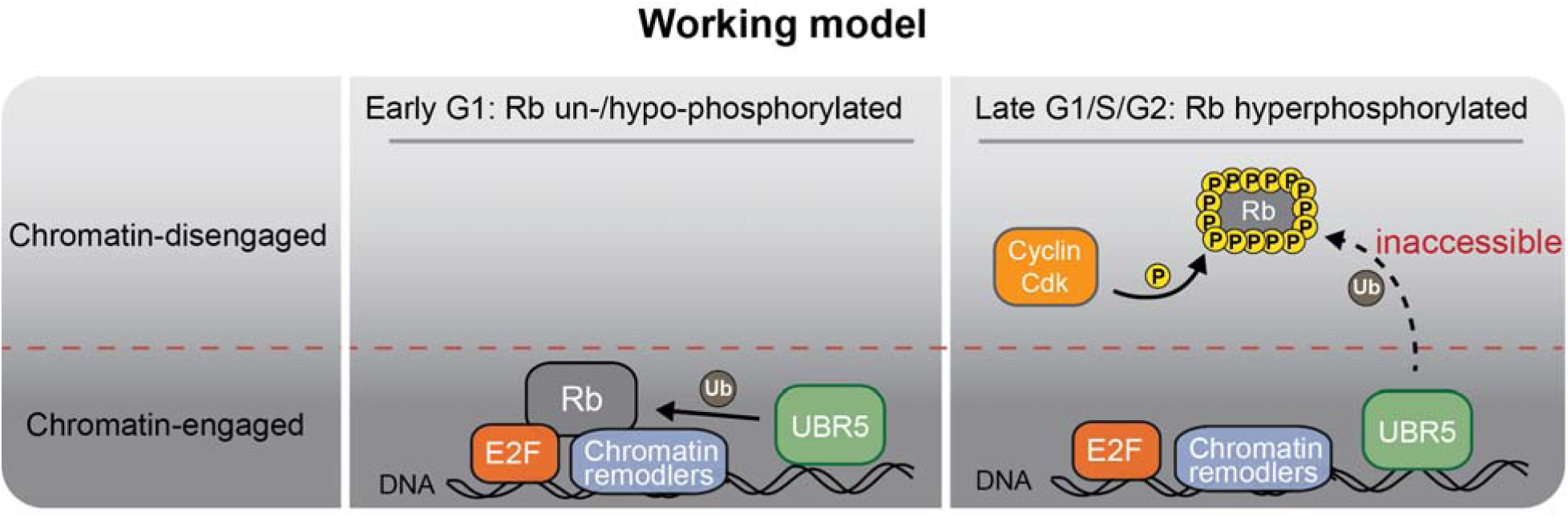
Model schematic for how chromatin association modulates Rb stability through the cell cycle. In early G1 phase, un-/hypo-phosphorylated Rb is associated with chromatin in complex with E2F-DP and chromatin remodelers and is targeted by UBR5 for degradation. In late G1, hyper-phosphorylation disrupts these interactions, promoting Rb dissociation from chromatin and rendering it inaccessible to UBR5-mediated degradation.

Our Rb chromatin association model is consistent with our observation that nearly all phosphorylation sites, except for the 3 N-terminal sites, contribute to Rb stability, with no single site exerting a dominant effect (Fig. 1B). Prior studies ^1–5,8,13–15^ found that phosphorylation at distinct sites disrupt different interaction interfaces between Rb and its binding partners, including the E2F-DP1 dimer and LxCxE containing proteins. Accordingly, Rb variants harboring different numbers of phospho-site mutations are expected to exhibit graded disruption of these interfaces, leading to corresponding differences in chromatin association and, consequently, different protein half-lives, as we observed. This framework explains why phosphomimetic Rb mutants display a cumulative increase in Rb half-life during early G1 (Fig. S1A). It may also account for the increased stability of both RbΔCDK variants bearing either an E-rich or Q-rich C-terminal tail (Fig. S1C). Because the Rb C-terminus is critical for its interactions with E2F-DP^3^, adding a long C-terminal tail may affect this interaction to reduce chromatin association and thereby stabilize these Rb variants.

While our data identify chromatin association as a key determinant of UBR5-mediated Rb degradation, it is unlikely to be the sole mechanism, because the RbΔCDK variants with disrupted interacting interfaces never reach the high stability observed for hyper-phosphorylated Rb (the wild-type Rb in S/G2 phase)^10^. Phosphorylation can induce extensive structural rearrangements of Rb^1–5,8^, raising the possibility that it also alters the surface by which Rb is recognized by UBR5. Addressing this possibility will require structural studies of the Rb-UBR5 interaction to define how unphosphorylated Rb is targeted and how phosphorylation disrupts this process. Moreover, phosphorylation not only reduces Rb chromatin association but also redistributes it to other chromatin loci, potentially through more transient interactions, as demonstrated by a recent ChIP-Seq study^30^. How this redistribution influences Rb stability remains unclear and will require further loci-specific molecular engineering. In addition, fusion of histone H1 modestly increased the stability of RbΔCDK (Fig. 5F; Fig. S11E). This effect is expected, as H1 variants themselves have exceptionally long half-lives (Fig. S11B), and their fusions could confer a general stabilizing influence on the fusion protein. Importantly, because our conclusions are based on relative comparisons between matched Rb variants with identical modifications (with or without H1 fusion), this overall stabilization does not affect our interpretations.

In conclusion, our study shows that CDK-dependent phosphorylation stabilizes Rb, at least in part, by dissociating it from chromatin and thereby reducing the probability of interaction with UBR5. This mechanism contrasts with other G1/S-regulated substrates, such as human p27 or yeast Sic1, where CDK phosphorylation directly generates phospho-degrons recognized by SCF ubiquitin ligases to promote degradation^48,49^. Instead, our results support a broader localization-based regulatory principle observed across many pathways, in which protein turnover is dictated by whether the substrate resides in the same subcellular compartment as a specific E3 ubiquitin ligase. For example, p53 and β-catenin degradation are gated by regulated nuclear localization^50^. While these examples highlight organelle level effects of localization on protein degradation, we find that this principle can also operate at a sub-organellar level. Here, Rb phosphorylation likely stabilizes Rb by shifting it from a chromatin-engaged, UBR5-accessible state to a chromatin-disengaged, but still nuclear, UBR5-inaccessible state. Thus, phosphorylation can control when and where ubiquitin-mediated protein degradation occurs not only through phospho-degrons, or altering organelle-level localization, but also through sub-organellar partitioning.

## Acknowledgements

The single molecule tracking (SMT) experiment described was supported, in part, by Award Number 1S10OD034400-01A1 from the S10 Instrumentation Programs. Its contents are solely the responsibility of the authors and do not necessarily represent the official views of the S10 Instrumentation Programs or the National Institutes of Health. We thank all members of the Skotheim laboratory for valuable discussions and feedback on this project. We thank Dr. Peter Pryciak for suggestions on experimental design. This work was supported by the NIH (4R00GM147351 grant to S.Z. and P01 CA254867 to J.M.S.)

## Author contributions

S.Z. conceived this study. S.Z., M.C.L., and J.M.S. designed the experiments. S.Z. and M.C.L. performed the experiments. S.Z., M.C.L., and J.K. analyzed the data. S.Z. and J.M.S. wrote the manuscript. S.Z., M.C.L., J.K., and J.M.S. edited the manuscript.

## Materials and Methods

### Cell culture conditions and cell lines

All cells were cultured at 37°C with 5% CO_2_. Non-transformed hTERT1-immortalized human mammary epithelium cells (HMEC) were obtained from Stephen Elledge’s laboratory at Harvard Medical School^51^ and cultured in MEGM™ Mammary Epithelial Cell Growth Medium (Lonza CC-3150). In microscopy experiments we used the same media but without phenol red to reduce background fluorescence (Lonza CC-3153 phenol-red free basal media supplemented with growth factors and other components from the Lonza CC4136 kit).

### Fluorescent reporter cell lines

The S/G2 FUCCI cell cycle reporter mCherry-Geminin^52^, the HDHB based CDK sensor^44^, and Rb C-terminus based CDK sensor^36^ were transduced into HMEC cells using lentivirus. The reporter plasmids, lentiviral packaging vector dr8.74, and the envelope vector VSVg were transfected into HEK 293T cells by PEI (1mg/mL, Sigma-Aldrich). 48 hours later, the lentivirus-containing medium was collected and used to infect HMEC cells. 2-3 days after infection, positive cells were sorted by FACS and expanded. The endogenously tagged *RB1-3xFLAG-Clover-sfGFP* HMEC cell line was created by Evgeny Zatulovskiy^53^.

### Inducible expression of Rb variants in cell lines

To inducibly express Rb variants in cells, we used a doxycycline-inducible Rb cassette ^35^, and performed site-directed mutagenesis (E0554S, NEB) to generate Rb variant plasmids. All the plasmids were PiggyBac transposon vectors containing the *RB1* variants fused with fluorescent Clover and 3xFLAG affinity tag sequences, a zeocin resistance gene, and a Tet-On 3G transactivator gene driven by the EF1α promoter. The amino acid sequence for the E-rich tail is: SSASNSNGEESGGLSGNNEEANGNNVGNEESSGSGGSVEEGSSGNSSGEESNNGSNNNEEGSGASGSNEEGGNGSGNNEENNSGGSSSEEGNNNNISAEENGNSNNNIEEGGVGSGNGEENNSSNLVGEESSNSGGGGEENSSNSGSGEEGSGGGSG; and the amino acid sequence for the corresponding Q-rich tail is: SSASNSNGQQSGGLSGNNQQANGNNVGNQQSSGSGGSVQQGSSGNSSGQQSNNGSNNNQQGSGASGSNQQGGNGSGNNQQNNSGGSSSQQGNNNNISAQQNGNSNNNIQQGGVGSGNGQQNNSSNLVGQQSSNSGGGGQQNSSNSGSGQQGSGGGSG.

The HMEC cell lines stably expressing doxycycline-inducible Rb variants were generated by transfecting cells with 1 µg of PiggyBac transposon vectors and 1 µg of PiggyBac Transposase plasmid using the FuGene HD reagent (Promega E2311). Zeocin (300 µg/ml) selection began two days after transfection and lasted for at least 2 weeks until all the cells became resistant.

### Live cell imaging and analysis

For half-life measurements using live-cell imaging, the cells were seeded on 35-mm glass-bottom dishes (CellVis) with 1 µg/ml Doxycycline added into the media. After two days of dox induction, dox containing media was withdrawn and cells were washed with PBS for 4 times, and replaced with phenol red free culture media. Then, the cells were transferred to a Zeiss Axio Observer Z1 microscope equipped with an incubation chamber and imaged for 48 hours at 37°C and 5% CO_2_. Brightfield and fluorescence images were collected at multiple positions every 20 minutes using an automated stage controlled by the Micro-Manager software. We used a Zyla 5.5 sCMOS camera and an A-plan 10x/0.25NA Ph1 objective. Since RbΔCDK is quickly degraded, the half-life analysis started 2 hours after doxycycline removal to ensure accurate quantification. The cell cycle phase was classified using an mCherry-Geminin FUCCI sensor. The early G1 phase traces were taken as those having no mCherry-Geminin expression that lasted longer than 7 hours. The S/G2 phase is defined by mCherry-Geminin FUCCI marker expression. For cells expressing the CDK sensor, the transition from low CDK activity to high CDK activity was taken as the inflection point of the cytoplasm-to-nuclear fluorescence ratio^44^.

### Fluorescence Recovery After Photobleaching (FRAP)

FRAP experiments were performed on a Zeiss LSM780 confocal microscope equipped with a 63x oil-immersion objective. The cells were seeded on Nunc Lab-Tek II 4-well chambered cover glass system (Thermo Scientific) and treated with 1 µg/ml Doxycycline for 48 hours before FRAP assay. These cells also expressed a CDK translocation sensor, which localizes to the nucleus under conditions of low CDK activity. For FRAP analysis, only cells displaying clear nuclear localization of the sensor were selected. A defined region of interest (ROI) within the nucleus was photobleached using high-intensity laser illumination, and fluorescence recovery was monitored over time at low laser power to minimize additional photobleaching. A second, unbleached nuclear ROI was used as a reference, and a background ROI outside the cell was used to correct for background fluorescence. Fluorescence intensities from both the bleach and reference ROIs were first background-subtracted. The fluorescence intensity of the bleached ROI was then normalized to that of the reference ROI at each time point to correct for overall photobleaching during image acquisition. The signal recovery half-time (t½) was calculated by fitting the fluorescence recovery curves to a single-exponential association model:

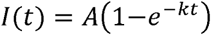

where *I(t)* is the normalized fluorescence intensity at time *t*, *A* is the mobile fraction, *1-A* is the immobile fraction, and *k* is the rate constant. The half-time of recovery was calculated as *t*_1/2_ = ln (2)/*k*.

### Single molecule tracking (SMT)

Cells were seeded on Nunc Lab-Tek II 4-well chambered cover glass (Thermo Scientific) and treated with 1 µg/mL doxycycline for 48 h prior to the SMT assay. Cells also expressed a CDK translocation sensor, which localizes to the nucleus under conditions of low CDK activity. For analysis, only cells displaying clear nuclear localization of the CDK sensor were selected. Photoactivatable Janelia Fluor 646 Haloalkane dye (Tocris, 8815) was added at a final concentration of 10 nM to label RbΔCDK variants. Cells were incubated with the dye at 37 °C for 30 min, followed by four rapid washes with warm, dye-free medium containing 0.1% BSA. Cells were then incubated in warm, dye-free medium for an additional 60 min, with two medium exchanges during this period. Fresh medium was added immediately before imaging.

Single-molecule imaging was performed using a Zeiss Elyra 7 microscope equipped with a 63x/NA 1.46 oil-immersion TIRF objective, a Hamamatsu ORCA Fusion BT camera, a Definite Focus 3 system, and a live-cell incubation chamber maintaining cells at 37 °C and 5% CO₂. The microscope was equipped with 405 nm (50 mW), 488 nm (500 mW), 561 nm (500 mW), and 640 nm (500 mW) laser lines. Highly inclined and laminated optical sheet (HILO) illumination was used for SMT. SMT experiments were performed using continuous low-power 405 nm illumination (0.1–5% laser power in ZEN Blue software) to photoactivate approximately 3–5 molecules per nucleus at any given time. Activated molecules were imaged using simultaneous fast-exposure illumination with a 640 nm laser (20% power) and an exposure time of 15 ms. A total of 2000 frames were acquired under continuous laser illumination at a camera frame interval of 18 ms. A single 488 nm image was acquired to determine CDK status and nuclear area.

Following image acquisition, single-particle tracking was performed using TrackMate in ImageJ. Molecules were localized in each frame using a Laplacian of Gaussian (LoG) detector with an estimated particle diameter of 0.8 µm. Trajectories were reconstructed by allowing a maximum displacement of 0.8 µm between consecutive frames and permitting a single-frame gap within a trajectory. Only particles localized within the nucleus were included in the analysis. Trajectories containing fewer than eight displacements were excluded from further analysis. Nuclear segmentation was performed using the CDK sensor image, as the sensor is nuclear under low-CDK conditions. The apparent diffusion coefficient (D) of individual molecules was calculated from the stepwise mean-squared displacement (MSD) of each trajectory according to:

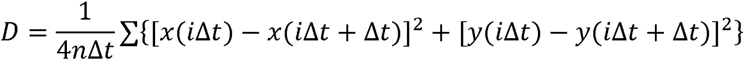

where *x(t)*and *y(t)*denote the coordinates of the molecule at time *t*, Δ*t* is the time interval between consecutive frames, and *n* is the number of steps in the trajectory.

To classify dynamic subpopulations, Gaussian mixture models (GMMs) were fit to the distributions of diffusion coefficients (*D*) across trajectories. Because both metrics are positively skewed, values were log-transformed prior to modeling. For each metric, univariate GMMs with *K* components were fit using maximum likelihood estimation, with *K* evaluated over a range (e.g., *K* == 2 - 3). The optimal number of components was selected by minimizing the Bayesian Information Criterion (BIC). *K* = 3 consistently yield the best fit for Rb variants, and K = 2 yield the best fit for Histone H1.2 and HaloTag-NLS. For each trajectory, posterior membership probabilities were computed from the fitted model, and trajectories were assigned to the most likely component (maximum posterior probability). The fraction of trajectories in each state was calculated from these assignments.

### Flow cytometry

For flow cytometry analysis, cells were grown on 6-well plates to ∼70% confluence and harvested following trypsinization. Cells were then stained with 20 µM Hoechst 33342 DNA dye in PBS for 30 minutes at 37°C and analyzed on an Attune NxT Flow Cytometer (Thermo Fisher Scientific). DNA content and the mCherry-Geminin fluorescent reporter were used to determine cell cycle phase, and the side scatter area parameter was used as a readout for cell size.

### RNA sequencing and data processing

Direct-zol RNA Miniprep kit (Zymo Research) was used to extract total RNA from 3 *UBR5+/+* clonal cell lines and 3 *UBR5-/-* clonal cell lines, which were generated from our previous study^10^. Two replicates were prepared for each clone. The total RNA was sent to Azenta for RNA-Seq sample processing, sequencing, and data analysis. Approximately 30 million paired-end reads were sequenced per sample, and TPM (Transcripts Per Million transcripts) were calculated for each sample (Supplementary table 1).

### TMT LC-MS/MS sample preparation

*UBR5+/+* and *UBR5-/-* cells (three clones for each background) were trypsinized, pelleted, and lysed for 30 minutes on ice in RIPA lysis buffer (Abcam) containing a protease and phosphatase inhibitor cocktail (Thermo). Cell lysates were cleared by centrifugation at 15000xg for 30 minutes at 4°C. The lysates were then denatured in 1% SDS, reduced with 5mM DTT, alkylated with 10mM iodoacetamide (Sigma), and then precipitated with three volumes of a solution containing 50% acetone and 50% ethanol. Proteins were re-solubilized in 2 M urea, 50 mM Tris-HCl, pH 8.0, and 150 mM NaCl, and then digested with TPCK-treated trypsin (50:1) overnight at 37°C. Trifluoroacetic acid and formic acid were added to the digested peptides for a final concentration of 0.2%. Peptides were desalted with a Sep-Pak 50mg C18 column (Waters). The C18 column was conditioned with 5 column volumes of 80% acetonitrile and 0.1% acetic acid and washed with 5 column volumes of 0.1% trifluoroacetic acid. After samples were loaded, the column was washed with 5 column volumes of 0.1% acetic acid followed by elution with 4 column volumes of 80% acetonitrile and 0.1% acetic acid. The elution was dried in a Concentrator at 45°C. Our method for TMT labeling was adapted from Zecha et al.^54^ and the Thermo TMT10plex™ Isobaric Label Reagent Set Protocol. In brief, acetone precipitated samples were resuspended in 100μm TEAB and digested O/N with TPCK trypsin (50:1) in the absence of Tris or Urea. After digestion, peptide concentration was ∼1μg/ul in 100μM TEAB for all samples. 20μg of peptide was labeled using 100μg of Thermo TMT10plex™ in a reaction volume of 25μl for 1 hour. The labeling reaction was quenched with 8μL of 5% hydroxylamine for 15 minutes. Labeled peptides were pooled, acidified to a pH of ∼2 using drops of 10% trifluoroacetic acid, and desalted with a Sep-Pak 50mg C18 column as described above.

### High-pH reverse phase fractionation

TMT-labeled peptides were fractionated using either an offline HPLC or the Pierce High pH Reversed-Phase Peptide Fractionation kit. HPLC fractionation was performed on *UBR5+/+* vs *UBR5-/-* dataset to achieve maximal depth (24 fractions). For IP experiments, the fractionation kit was used. The eight default fractions were used. In all cases, dried peptides were reconstituted in 0.1% TFA. Peptide concentrations were determined using a Nanodrop before injection.

### LC-MS/MS data acquisition

Desalted TMT-labeled peptides were analyzed on a Fusion Lumos mass spectrometer (Thermo Fisher Scientific, San Jose, CA) equipped with a Thermo EASY-nLC 1200 LC system (Thermo Fisher Scientific, San Jose, CA). Peptides were separated by capillary reverse phase chromatography on a 25 cm column (75 μm inner diameter, packed with 1.6 μm C18 resin, AUR2-25075C18A, Ionopticks, Victoria Australia). Electrospray Ionization voltage was set to 1550 volts. Peptides were introduced into the Fusion Lumos mass spectrometer using a two-step linear gradient with 6–33% buffer B (0.1% (v/v) formic acid in 80% acetonitrile) for 145 min followed by 33-45% buffer B for 15 min at a flow rate of 300 nL/min. Column temperature was maintained at 40°C throughout the procedure. Xcalibur software (Thermo Fisher Scientific) was used for the data acquisition, and the instrument was operated in data-dependent mode. Survey scans were acquired in the Orbitrap mass analyzer over the range of 380 to 1800 m/z with a mass resolution of 70,000 (at m/z 200). Ions with a charge state of either 2, 3 or 4 were selected for fragmentation within an isolation window of 0.7 m/z. Selected ions were fragmented by Higher-energy Collision-induced dissociation (CID) with normalized collision energies of 35% and the tandem mass spectra was acquired in the Ion trap mass analyzer with a with a “Rapid” scan rate. Repeated sequencing of peptides was kept to a minimum by dynamic exclusion of the sequenced peptides for 60 seconds. For MS/MS, the AGC target was set to “Standard” and max injection time was set to 35ms. Relative changes in peptide concentration were determined at the MS3-level by isolating and fragmenting the 9 most dominant MS2 ion peaks using HCD. TMT reporter ions were resolved in the orbitrap at a resolution of 50,000.

### Spectral searches

All raw files were searched using the Andromeda engine^55^ embedded in MaxQuant (v2)^56^. For TMT searches, a Reporter ion MS3 search was conducted using 10plex or 16plex TMT isobaric labels. Variable modifications included oxidation (M) and protein N-terminal acetylation. Carbamidomthyl (C) was a fixed modification. The number of modifications per peptide was capped at five. Digestion was set to tryptic (proline-blocked). Database search was conducted using the UniProt proteome - Human_UP000005640_9606. Minimum peptide length was 7 amino acids. FDR was determined using a reverse decoy proteome^57^.

### Two-dimensional annotation enrichment analysis

Annotation enrichment analysis was performed as described previously^58^. The protein annotation groups were deemed significantly enriched and plotted if the Benjamini–Hochberg FDR <0.02. The position of each annotation group on the plot is determined by the mean of Log_2_ ratio (KO/WT) of mRNA or protein.

### Subcellular compartment annotation analysis on post-transcriptionally regulated genes

First, mass spectrometry peptide data was cleaned and filtered for a minimum of 5 unique peptides per protein annotation. False Discovery Rates (FDR) were calculated for each protein using the Benjamini-Hochberg correction. Significantly changed proteins (FDR < 0.05) were selected for downstream clustering. Next, FDR results were calculated for TPM data from bulk mRNA sequencing. Proteins corresponding to insignificantly changed mRNAs were intersected with the significantly changed proteins to obtain all post-transcriptionally enriched proteins in *UBR5-/-* cells. Gene Ontology annotations for this set were obtained and used to identify corresponding subcellular compartments using SubcellulaRVis^18^.

### Immunoprecipitation of Rb variants

Rb complex extraction and immunoprecipitation (IP) was performed as in ^13^. Briefly, HMEC cells transduced with Dox-inducible Rb were treated with 1 μg/mL Dox for 48h to induce Rb variants protein. Palbociclib treatment (1µM) started 24h before harvest to synchronize cells in G1 phase. Approximately 10^7^ cells per culture were washed twice with ice-cold PBS and cell lysates were collected in 1 mL E1Agl^250^ buffer (50 mM HEPES-KOH pH 7.4, 250 mM NaCl, 0.1% NP-40, 10% glycerol, 1 mM PMSF) supplemented with Thermo protease and phosphatase inhibitors. Lysates were incubated on ice for 30 min and then were sonicated with 2×15s pulses (45 s interval) and amplitude 50%. Lysates were centrifuged at 16,000 x g for 15 min at 4°C and supernatants were incubated with anti-FLAG M2 magnetic beads under rotation for 2h at 4°C (50 μL of magnetic beads per IP). Beads were washed three times with 1 mL E1Agl^150^ buffer (50 mM HEPES-KOH pH 7.4, 150 mM NaCl, 0.1% NP-40, 10% glycerol, 1 mM PMSF) and resuspended with elution buffer (100 mM Tris-HCl (pH 8.0), 1 % SDS). Beads were incubated at 70°C for 10 min to elute. After separating elution with beads, the eluted proteins were reduced with 5mM DTT and prepared in the same way as the LC-MS/MS samples. An aliquot of elution was analyzed by western blot. For western blot analysis, 5% of input cell lysate was kept before IP. IP samples from cells lacking Dox-induced Rb expression were used as negative controls. Proteins identified with at least three unique peptides and exhibiting no less than 2-fold higher proportion in any Rb variant pull-down relative to the negative control were considered as Rb-interacting proteins. Normalized enrichment score were calculated by normalizing the proportion of each interacting protein to the corresponding bait Rb variant, with the RbΔCDK set to 1.

### Immunoblots

Cell lysates or eluted IP samples were separated on NuPAGE 3-8% Tris-Acetate protein gels (Thermo Fisher Scientific) and transferred to nitrocellulose membranes. Membranes were then blocked with SuperBlock™ (TBS) Blocking Buffer (Thermo Fisher Scientific) and incubated overnight at 4°C with primary antibodies in 3% BSA solution in PBS. The primary antibodies were detected using the fluorescently labeled secondary antibodies IRDye® 680LT Goat anti-Mouse IgG (LI-COR 926-68020) and IRDye® 800CW Goat anti-Rabbit IgG (LI-COR 926-32211). Membranes were imaged on a LI-COR Odyssey CLx and analyzed with LI-COR Image Studio software. The following primary antibodies were used: anti-Rb (Santa Cruz, sc-74570, 1:500), anti-HDAC1 (CST#5356, 1:1000).

### Image analysis

For live-cell imaging microscopy data, cell nuclei were segmented using the GFP channel from the inducibly overexpressed Clover-3xFlag-Rb variants. Segmentation was performed using the Fiji plugin StarDist, which is a deep-learning tool for segmenting nuclei in images that are difficult to segment using thresholding-based methods. The total pixel intensities within the segmented masks in each channel were recorded, and each object’s background was subtracted based on the median intensity of the image. The tracking of live cells was done manually using the TrackMate plugin in Fiji. To determine protein half-life, fluorescence intensity degradation traces were fitted to a single-exponential decay model*, N(t) == N_O_e^-kt^,* where *N(t)*is the fluorescence intensity at time *t, N_O_* is the initial intensity, and k is the degradation rate constant. Protein half-life (*t*_l/2_) was calculated as *t*_l/2_ == ln (2)/*k*.

## Statistical analysis

The data in most Fig. panels reflect multiple biological replicate experiments performed on different days. For comparison between groups, we generally conducted unpaired two-tailed Student’s t-tests. Statistical significance is displayed as p < 0.05 (∗) or p < 0.01 (∗∗) unless specified otherwise.

