## Supplemental figures for "Chromatin association promotes UBR5-mediated degradation of Rb"

**Figure S1**

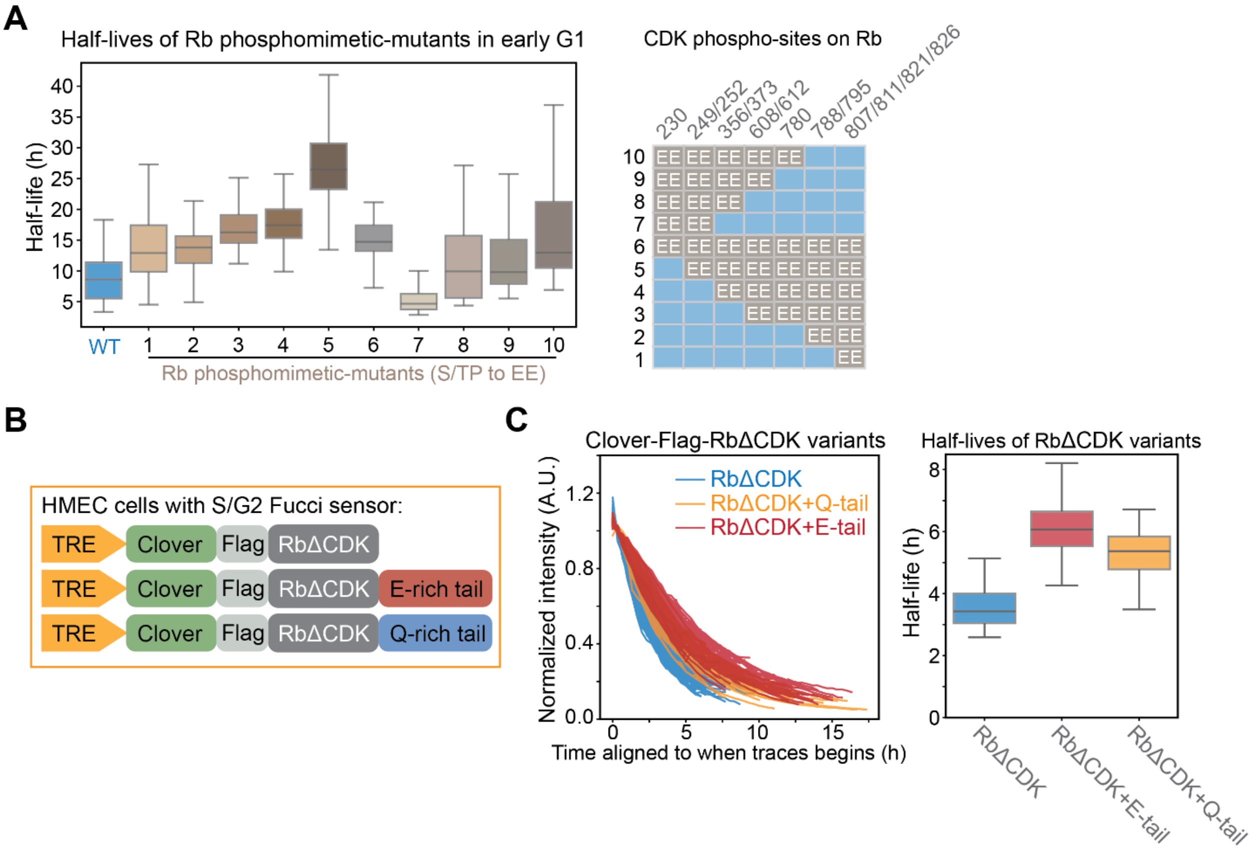


**Figure S1. Phosphorylation mediated Rb stabilization is not driven by charge-mediated mechanism.**

**A.** Left panel shows the half-life distribution of Rb phosphomimetic-mutants (S/TP to EE) in early G1 phase. The data are from our previous study[10]. Right panel shows the phosphorylation sites mutated in each Rb variant. **B.** Schematic illustrating the experimental strategy used to test whether the indicated mutations affect Rb protein stabilization. **C.** Left panel shows the degradation traces of Clover-3xFlag-RbΔCDK and variants carrying either a charge-mimicking E-rich tail or a control Q-rich tail following Dox withdrawal. Only the traces in early G1 phase were selected, and the classification was based on a FUCCI cell cycle marker and cell cycle phase duration. Right panel shows the distribution of half-lives estimated from exponential fits on the traces in left panel. Box plot indicates 5th, 25th, median, 75th, and 95th percentiles.

**Figure S2**

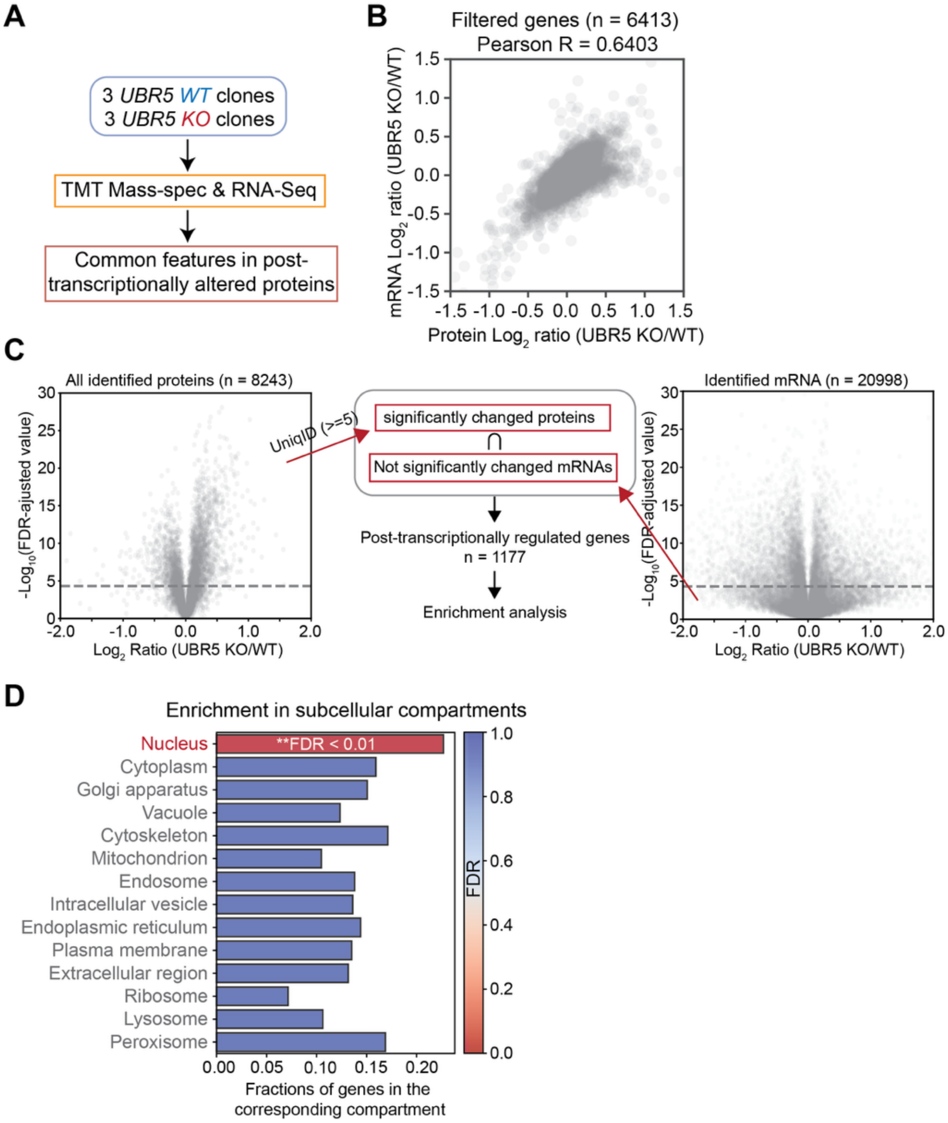


**Figure S2. UBR5 preferentially targets nucleus localized proteins.**
**A.** Workflow for RNA-Seq and proteomics analyses performed on *UBR5 WT* and *KO* clonal cell lines. **B.** Pearson correlation between the $\log_2$ transformed protein concentration ratios (*UBR5 KO*/*WT*; x-axis) and mRNA concentration ratios (*UBR5 KO*/*WT*; y-axis). **C.** Volcano plots showing the $\log_2$ transformed concentration ratios (KO/WT) for proteins identified by quantitative proteomics and transcripts identified by RNA-Seq. The middle panel illustrates the strategy used to identify post-transcriptionally regulated genes. **D.** Subcellular compartment enrichment analysis of genes showing post-transcriptionally regulation in *UBR5 KO* cells relative to *UBR5 WT* cells.

**Figure S3**

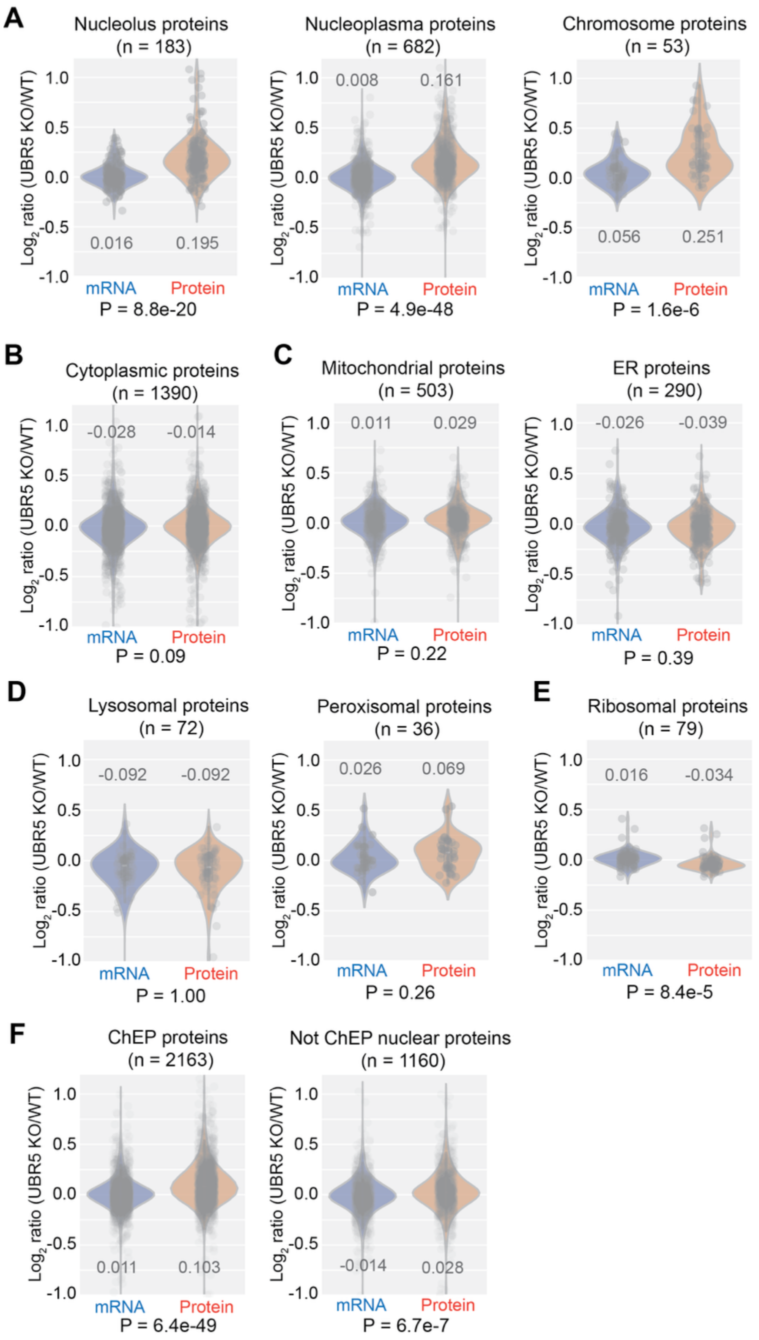


**Figure S3. UBR5 preferentially targets chromatin-associated proteins.**
**A-E.** Violin plots showing the distribution of $\log_2$ transformed mRNA and protein ratios (KO/WT) for genes grouped by subcellular localization. Subcellular localization was assigned based on UniProt annotations. Each dot represents one gene, and the total number of genes in each group is indicated in the plot title. The $\log_2$ (KO/WT) values and P values comparing protein and mRNA concentration changes are indicated on the plots. **F**. The $\text{Log}_2$ ratios (UBR5 KO/WT) in mRNA and protein levels for proteins identified in ChEP (Chromatin enrichment for proteomics) (left) and proteins not identified by ChEP but are nuclear (right). The proteins identified by ChEP are

defined in [29], and the non-ChEP nuclear proteins are defined using the data from the same study[29], where the proteins are classified as “Nuclear” but not identified by ChEP. Each dot represents one gene, and the total number of genes is indicated in the plot title. The $Log_2$ (UBR5 KO/WT) values and P-value comparing protein and mRNA concentration changes is indicated on the plot.

**Figure S4**

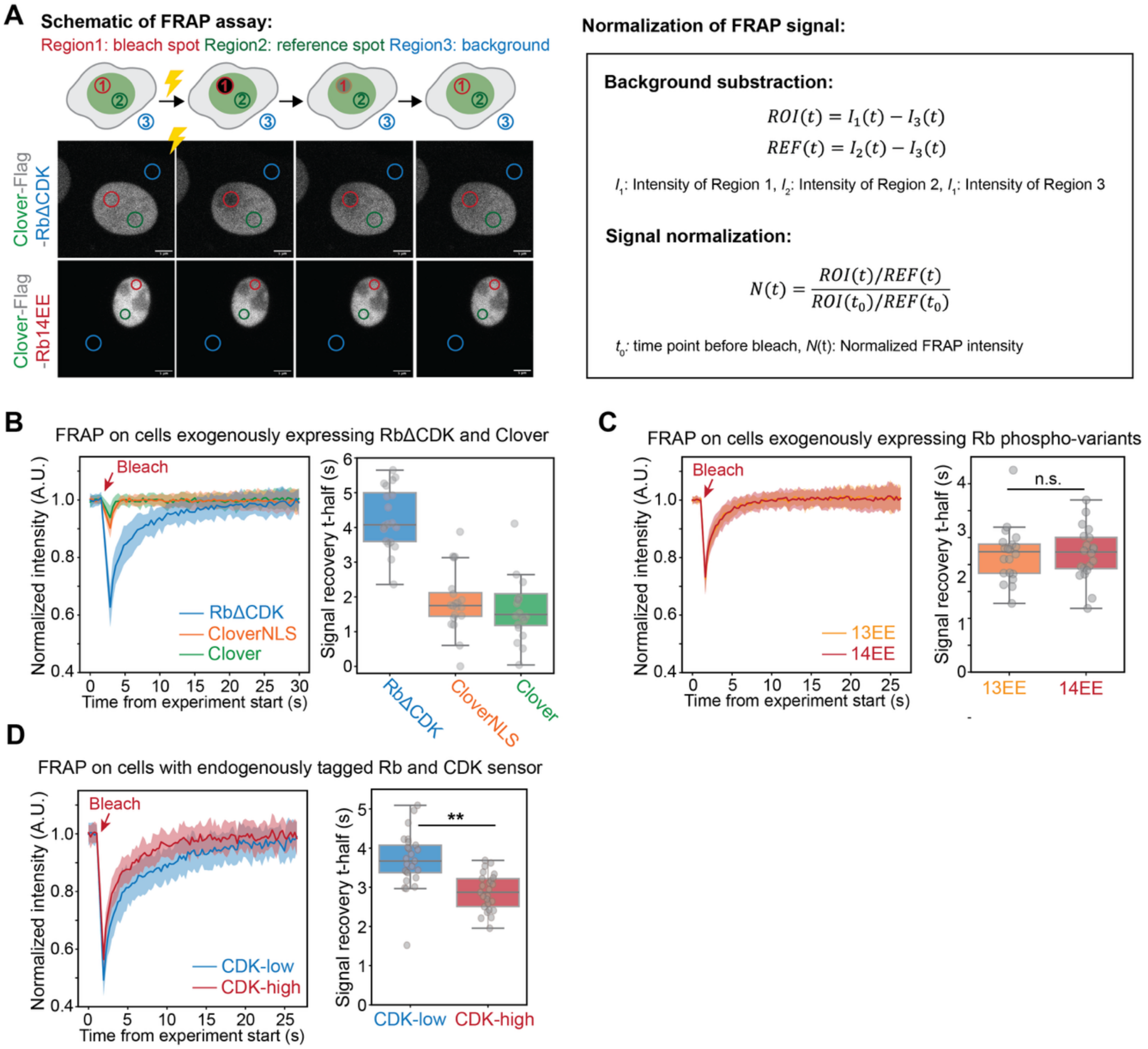


**Figure S4. Rb phosphorylation reduces its chromatin association.**
**A.** Left panel shows the schematic of the FRAP assay. Representative cells expressing Clover-3xFlag-RbΔCDK or Clover-3xFlag-Rb14EE Rb protein variants are shown. Right panel shows the normalization procedure for FRAP signal, using the intensities obtained from Region1, 2, and 3. **B.** Left panel shows the mean normalized FRAP recovery curve from HMEC cells expressing Clover-3xFlag-RbΔCDK, CloverNLS, or Clover. The bleach time point is indicated; shaded area denotes standard deviation. Right panel shows the distribution of FRAP signal recovery half-time (t-half) for RbΔCDK, CloverNLS, and Clover. Box plot shows the 5$^{th}$, 25$^{th}$, median, 75$^{th}$, and 95$^{th}$ percentiles. Each dot represents one measured cell. **C.** Left panel shows the mean normalized FRAP recovery curve from HMEC cells expressing Clover-3xFlag-Rb13EE or Clover-3xFlag-Rb14EE. Only CDK-low cells were selected for experiment. The bleach time point is indicated; shaded area denotes standard deviation. Right panel shows the distribution of FRAP signal recovery half-time (t-half) for Rb13EE and Rb14EE protein variants. Box plot shows the 5$^{th}$, 25$^{th}$, median, 75$^{th}$, and 95$^{th}$ percentiles. Each dot represents one measured cell. There is no significant difference between the two Rb variants. **D.** Left panel shows the mean normalized FRAP recovery curves from HMEC cells expressing endogenously

tagged *RB1-3xFlag-Clover* and the CDK sensor[36]. The bleach time point is indicated; shaded area denotes standard deviation. Right panel shows the distribution of FRAP signal recovery half-time (t-half) for Rb in CDK-low and CDK-high states. Box plot shows the 5th, 25th, median, 75th, and 95th percentiles. Each dot represents one measured cell. ***P*<0.01.

**Figure S5**

**A**

**Specific mutations for each Rb mutant type**

| Name | Amino acid change |
|---|---|
| dG | R467A, K548A |
| LCE | I753A, N757A, M761A |
| C7A | R828A,S829A,I831A,V833A,I835A,F845A,N849A |

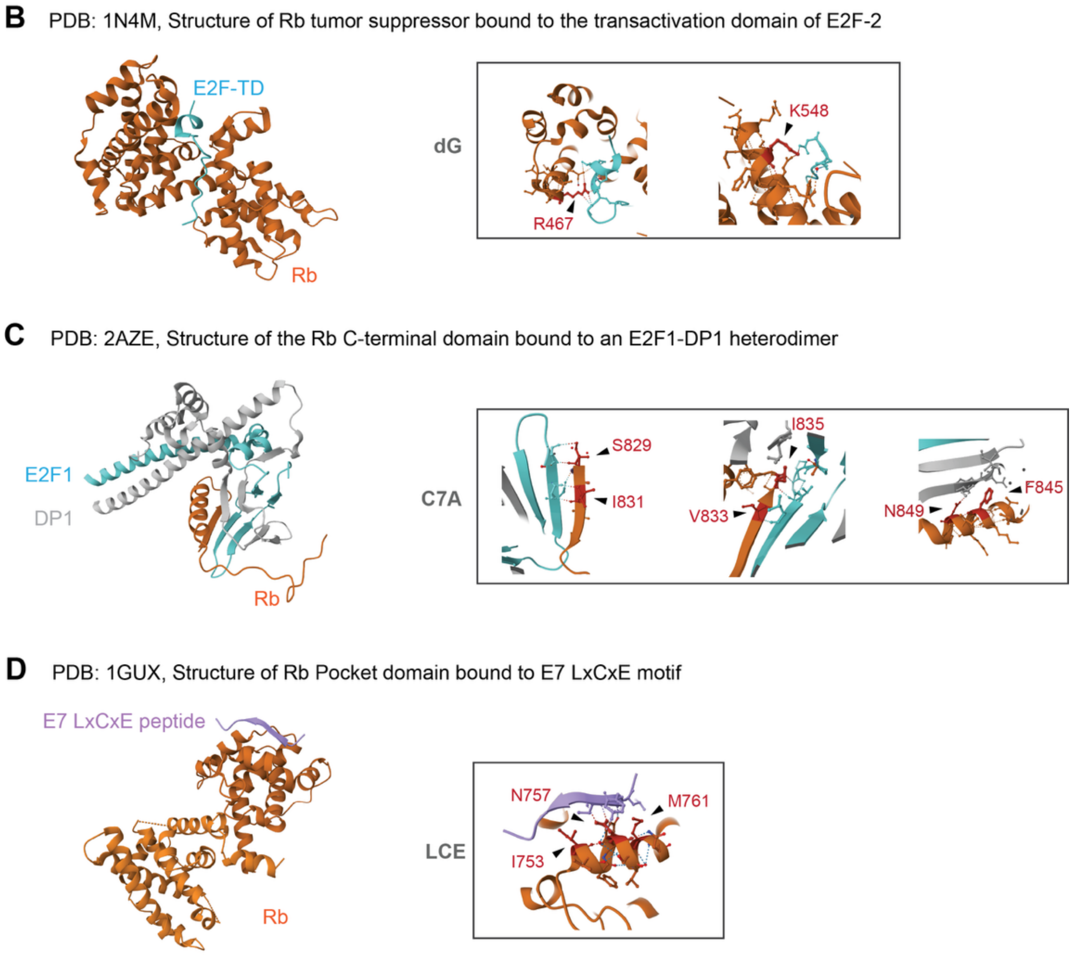


**Figure S5. Previously solved crystal structures of Rb's interactions with E2F, DP1, and E7.** **A.** The amino acid substitutions corresponding to the indicated Rb variant. **B.** Left panel shows the crystal structure of Rb pocket domain bound to E2F2-TD (PDB: 1N4M). Right panel shows the magnified view highlighting the mutated residues in the dG Rb mutants. **C.** Left panel shows the crystal structure of Rb C-terminal domain bound to an E2F1-DP1 dimer (PDB: 2AZE). Right panel shows the magnified view highlighting the mutated residues in the C7A Rb mutants. **D.** Left panel shows the crystal structure of the Rb pocket domain bound to the E7 LxCxE motif (PDB: 1GUX). Right panel shows the magnified view highlighting the mutated residues in the LCE Rb mutant.

**Figure S6**

**A**

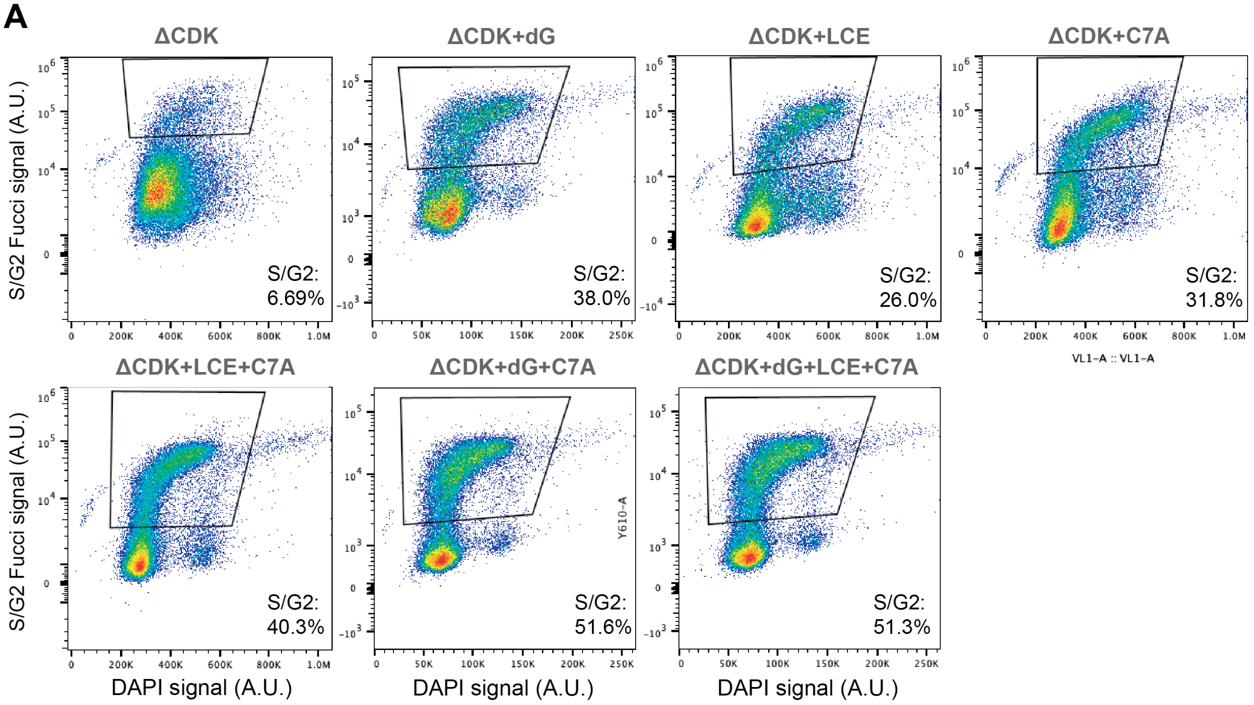


**Figure S6. Cell cycle profiles of the RbΔCDK variants.**

**A.** Flow cytometry analysis of HMEC cells expressing Clover-3xFlag-RbΔCDK variants together with a FUCCI S/G2 cell cycle marker. DNA content (DAPI) is shown on the x-axis, and FUCCI S/G2 reporter signal is shown on the y-axis. The boxed region indicates the S/G2 cells as defined by the FUCCI marker. The percentages of S/G2 cells are indicated in each plot.

**Figure S7**

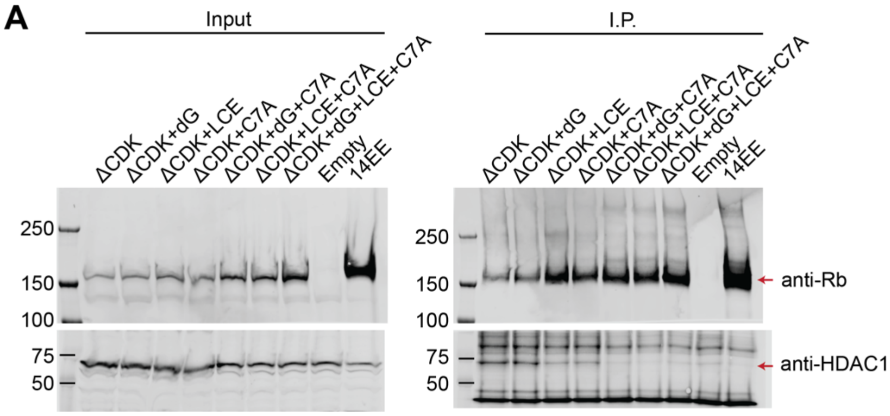


**Figure S7. IP-MS analysis of the interactome of Rb variants.**
**A**. HMEC cells expressing Clover-3xFlag-Rb variants were induced with 1µg/ml Dox for 48 hours and treated with Palbo (1µM) for 24 hours before collection. Clover-3xFlag-Rb variants were immunoprecipitated using anti-Flag magnetic beads. The samples were processed for mass spectrometry and also analyzed by immnunoblotting for Rb and HDAC1.

### Figure S8

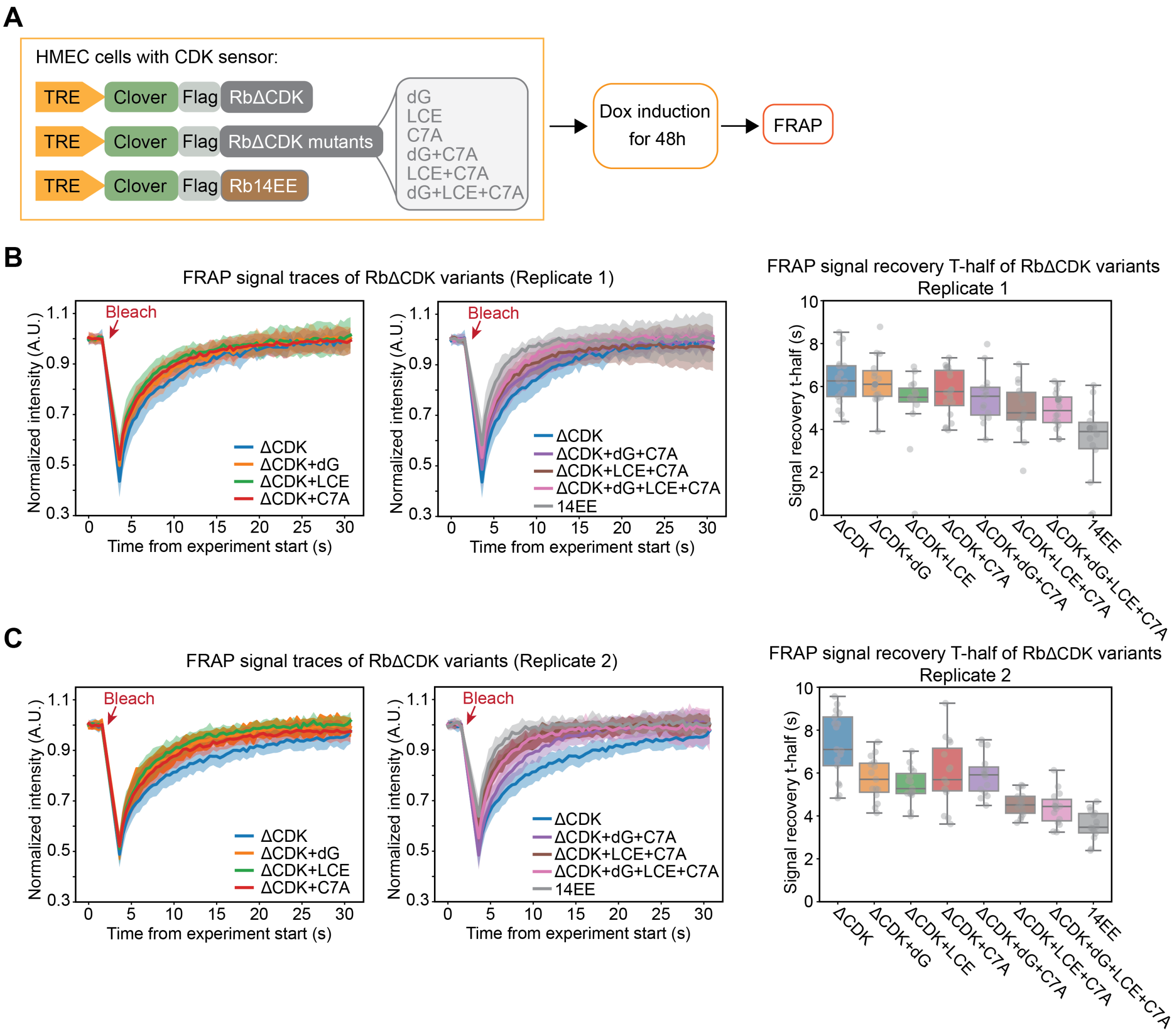


**Figure S8. Rb variants exhibit reduced chromatin association as measured by FRAP. A.** Schematic illustrating the cell lines expressing different Rb variants and the experimental workflow for the FRAP assay. **B-C.** Left panels shows the mean normalized FRAP recovery curves from HMEC cells expressing Clover-3xFlag-Rb variants for replicate 1 (**B**) and replicate 2 (**C**). The bleach time point is indicated; shaded area denotes standard deviation. Right panels show the distribution of FRAP signal recovery half-time (t-half) for the indicated Rb variants for replicate 1 (**B**) and replicate 2 (**C**). Box plot shows the $5^{th}$, $25^{th}$, median, $75^{th}$, and $95^{th}$ percentiles. Each dot represents one cell.

**Figure S9**

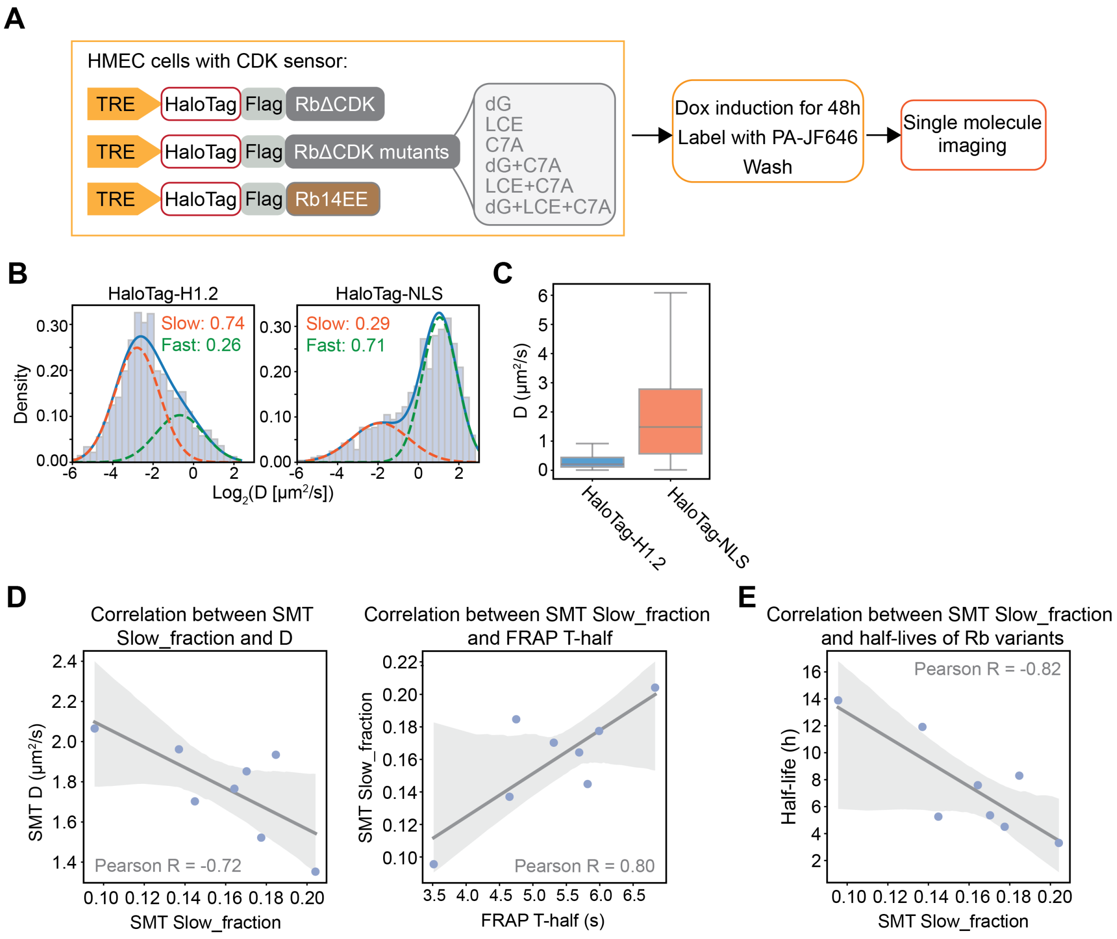


**Figure S9. Rb variants exhibit reduced chromatin association as measured by single molecule tracking (SMT).**
**A.** Schematic illustrating the cell lines expressing different Rb variants and the experimental workflow for the SMT assay. **B.** Probability densities of $log_2$-transformed apparent diffusion coefficients ($Log_2$ D) for HaloTag fused histone H1.2 or HaloTag-NLS, measured by SMT. The distribution was fit using a Gaussian mixture model (GMM). The solid blue line represents the overall GMM probability density function (PDF), while dotted lines indicate the PDFs of individual mixture components. The fraction of molecules assigned to each diffusive state (Slow, Fast) was estimated from the fitted mixture weights and is indicated on the plot. **C.** Box plot of the diffusion coefficient (D) for HaloTag fused histone H1.2 or HaloTag-NLS measured by SMT. Box plots denote the first quartile, median, and third quartile. **D.** Left panel shows the Pearson correlation between D and slow fraction measured from SMT across Rb variants. Right panel shows the Pearson correlation between the slow fraction measured from SMT and the FRAP recovery half-time (t-half) across Rb variants. Shaded area denotes the 95% confidence interval of a linear fit. **E.** Pearson correlation between the slow fraction measured from SMT and the protein half-lives across Rb variants. Shaded area denotes the 95% confidence interval of a linear fit.

**Figure S10**

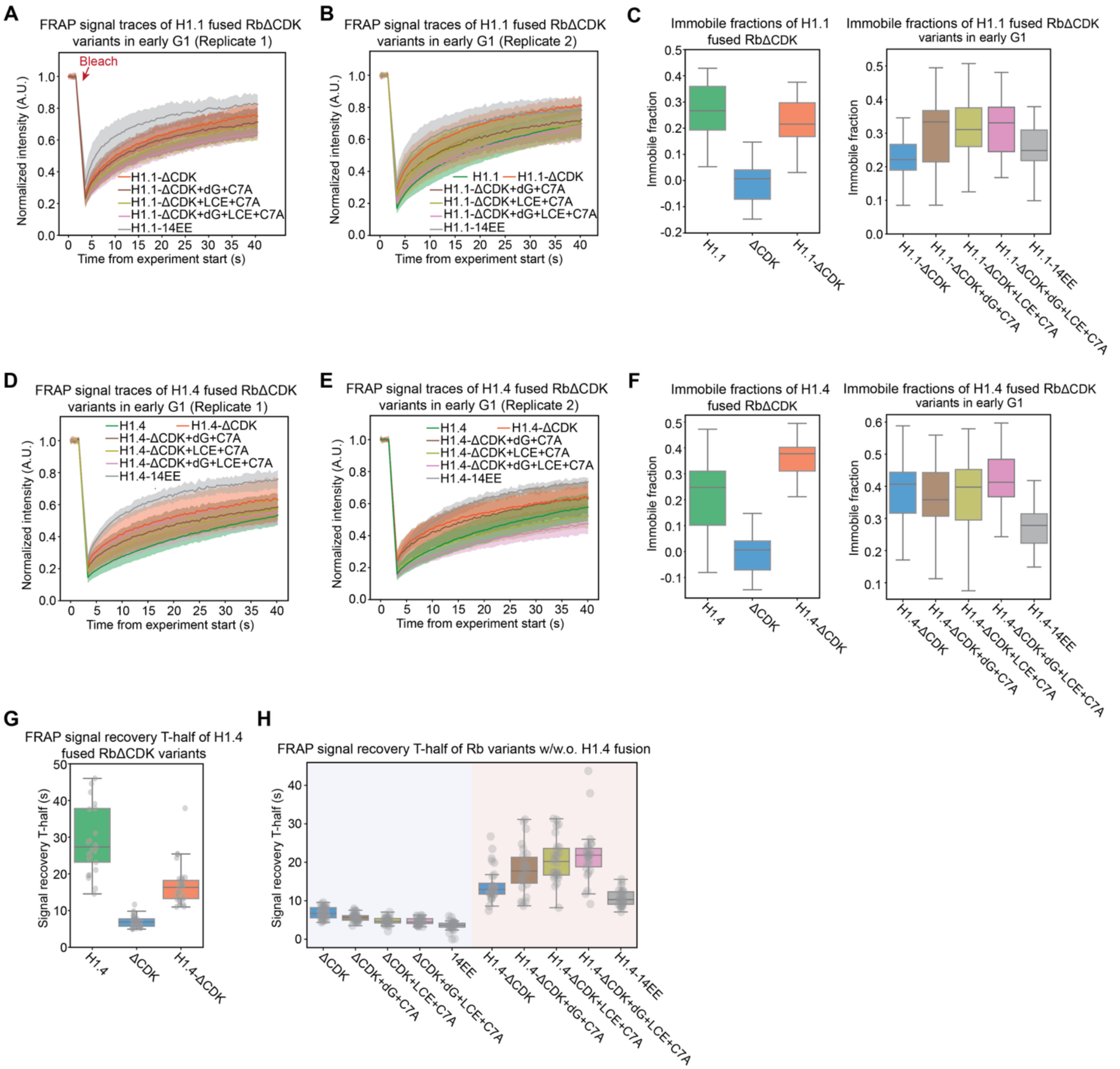


**Figure S10. Histone H1 fusion increase chromatin association of Rb variants.**
**A-B**. Mean normalized FRAP recovery curves from HMEC cells expressing H1.1 fused Clover-3xFlag-Rb variants for replicate 1 (**A**) and replicate 2 (**B**). **C.** Distribution of immobile fractions measured by FRAP for the indicated H1.1 fused Rb variants. Box plot shows the $5^{th}$, $25^{th}$, median, $75^{th}$, and $95^{th}$ percentiles. Each dot represents one cell. **D-E**. Mean normalized FRAP recovery curves from HMEC cells expressing H1.4 fused Clover-3xFlag-Rb variants for replicate 1 (**C**) and replicate 2 (**D**). **F.** Distribution of immobile fractions measured by FRAP for the indicated H1.4 fused Rb variants. Box plot shows the $5^{th}$, $25^{th}$, median, $75^{th}$, and $95^{th}$ percentiles. Each dot represents one cell. **G.** Distribution of FRAP signal recovery half-time (t-half) for the indicated Rb variants. Box plot shows the $5^{th}$, $25^{th}$, median, $75^{th}$, and $95^{th}$ percentiles. Each dot represents one cell. **H.** Distribution of FRAP signal recovery half-time (t-half) for the indicated

Rb variants with or without H1.4 fusions. Box plot shows the $5^{th}$, $25^{th}$, median, $75^{th}$, and $95^{th}$ percentiles. Each dot represents one cell.

**Figure S11**

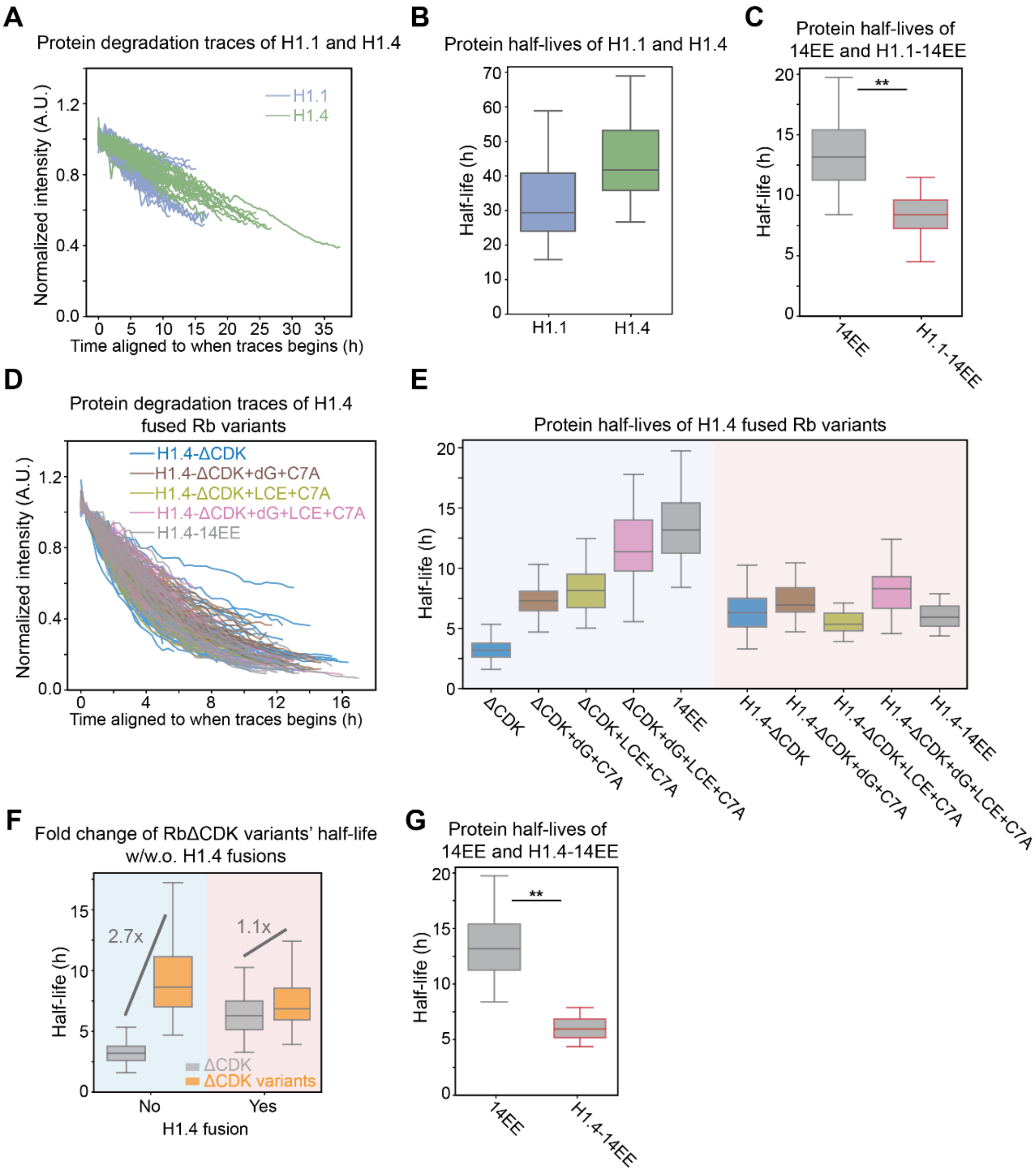


**Figure S11. Tethering Rb variants to chromatin prevents their stabilization.**
**A.** Degradation traces of H1.1-Clover and H1.4-Clover following Dox withdrawal. Only the traces in early G1 phase were selected, and the classification was based on a FUCCI cell cycle marker and cell cycle phase duration. **B.** Distribution of half-lives estimated from exponential fits on the traces in (**A**). Box plot indicates 5$^{th}$, 25$^{th}$, median, 75$^{th}$, and 95$^{th}$ percentiles. **C.** Distribution of half-lives estimated from exponential fits on the traces in **Fig. 5E**. Box plot indicates 5$^{th}$, 25$^{th}$, median, 75$^{th}$, and 95$^{th}$ percentiles. ** *P*<0.01. **D.** Degradation traces of H1.4-fused Clover-3xFlag-Rb variants following Dox withdrawal. Only the traces in early G1 phase were selected, and the classification was based on a FUCCI cell cycle marker and cell cycle phase duration. **E.** Distribution of half-lives estimated by exponential fitting of the traces in (**D**), compared with the corresponding non-H1.4-fused Rb variants (**Fig. 4C**). Box plot indicates 5th, 25th, median, 75$^{th}$,

and 95th percentiles. **F.** Half-life distribution for RbΔCDK (grey) and pooled RbΔCDK variants carrying two or three mutation sets (orange) with or without H1.4 fusion. Variant half-lives were pooled together from dG-C7A, LCE-C7A, and dG-LCE-C7A in (**E**). The fold-changes relative to RbΔCDK are indicated in the plot. **G.** Distribution of half-lives estimated from exponential fits on the traces in **(E)**. Box plot indicates 5$^{th}$, 25$^{th}$, median, 75$^{th}$, and 95$^{th}$ percentiles. ** *P*<0.01.
